# Single-cell quantification of the iron-neuromelanin balance in dopaminergic neurons across the lifespan

**DOI:** 10.64898/2026.05.08.721830

**Authors:** Felix Büttner, Tilo Reinert, Carsten Jäger, Malte Brammerloh, Markus Morawski, Ilona Lipp, Gerald Falkenberg, Dennis Brückner, Richard McElreath, Catherine Crockford, Roman Wittig, Tobias Deschner, EBC Consortium, Nikolaus Weiskopf, Evgeniya Kirilina

## Abstract

Dopaminergic neurons in the substantia nigra depend on iron for dopamine synthesis but are vulnerable to iron-induced oxidative stress. Many of these neurons synthesize neuromelanin, an iron-chelating pigment that accumulates across the lifespan and makes them vulnerable in Parkinson’s disease. It remains unclear whether their selective vulnerability arises from neuromelanin overload or from the release of toxic labile iron from the oversaturated pigment. We quantified iron and neuromelanin at the single-cell level across the lifespan of chimpanzees, a species closely resembling humans in pigment and iron accumulation. Combining quantitative MRI, X-ray fluorescence imaging, and microscopic colorimetry, we found that the iron-to-neuromelanin ratio remains stable with age across large neuronal populations. Chemical equilibrium modeling of the iron binding in neuromelanin indicated that cytosolic labile iron concentrations remain low throughout adulthood. We have found no evidence for neuromelanin saturation or increased iron-mediated toxicity with age. This finding challenges the hypothesis that neuromelanin saturation drives age-related dopaminergic vulnerability. The presented method provides a quantitative framework for studying iron homeostasis in these neurons.

## 1 Introduction

The depletion of iron-rich, neuromelanin-pigmented dopaminergic neurons (DN) in the substantia nigra (SN) is a hallmark of Parkinson’s disease [1]. Despite being recognized for more than a century [2, 3], the basis of the selective vulnerability of these pigmented neurons remains unresolved [4–6]. One prominent hypothesis linked their susceptibility to the lifespan accumulation of neuromelanin (NM), a metal-chelating biopolymer formed from reactive byproducts of dopamine metabolism, which may impair cellular function if occupying most of the cytosol [7–11]. An alternative hypothesis posits that NM becomes increasingly saturated with iron over time and may begin to release poorly bound iron into the cytosol, thereby driving oxidative stress and neurodegeneration [6, 8, 12–16].

Iron is essential for the dopamine synthesis, forming the catalytic center of the dopamine-regulating enzyme tyrosine hydroxylase (TH). Thus, DN require sufficient iron levels for normal function [11, 13, 17]. At the same time, labile iron within the cellular cytosol is an important driver of cellular vulnerability, inducing oxidative stress, triggering iron-induced cell death (ferroptosis) and *α*-synuclein aggregation [13, 18–20]. Moreover, in DN, labile iron promotes the oxidation of dopamine to toxic quinone species, and therefore dopamine and iron have been referred to as a “toxic couple” [6, 17, 19, 21]. The brown pigment NM, composed of pheomelanin and eumelanin monomers, was originally considered a neuroprotective chelator that shields neurons from oxidative stress, reactive dopamine quinones, and ferroptosis [17, 22]. With age, however, some DN accumulate large amounts of NM, which may either occupy a substantial fraction of the cytoplasm and impair cellular function [7, 8, 11, 23, 24] or become progressively saturated with iron, thereby losing its chelating capacity and releasing labile redox-active iron that exacerbates oxidative stress [6, 13, 15, 16].

Yet, the relative contribution of NM accumulation versus increasing iron toxicity to age-related dopaminergic neuronal vulnerability remains unknown, a question of particular relevance given the recent emergence of iron-chelating therapies [25, 26]. The main reason for this knowledge gap is the absence of methods – invasive or non-invasive – that can simultaneously quantify iron and NM accumulation at the level of individual cells across the lifespan.

Recent advances in quantitative MRI (qMRI) offer promising non-invasive markers of tissue iron and potentially NM, linking the transverse relaxation rates and other qMRI parameters to iron and NM content [27–34]. However, due to limited spatial resolution, these MRI-based measures yield only macroscopic tissue averages and it remains unclear which cells and other tissue components contribute to the MRI signal. Also histochemically, lifespan NM and iron accumulation have so far been assessed only as bulk tissue concentrations from *post mortem* human brain extracts [6, 8, 13, 14, 35] providing limited insights into the cell-specific iron levels of DN, which differ from other cells [36]. Homogenization and extraction of NM changes the iron loading of the iron-binding sites, making bulk iron measurements poorly suited to studying the physiological iron saturation of NM [37, 38]. Recent progress in physical methods for tissue iron quantification has enabled accurate measurements of cellular iron loads in *post mortem* sections with micrometer resolution using X-ray fluorescence imaging (μXRF) [39]. So far, these techniques have only been applied to a small number of cells mostly from elderly individuals, and have not been used to systematically characterize iron accumulation across the entire lifespan [15, 27, 36, 40–44].

Another major bottleneck is the limited availability of *post mortem* tissue from young human individuals and the lack of appropriate animal models with human-like NM and iron accumulation throughout the lifespan. Common laboratory rodents naturally lack NM and exhibit very low brain iron levels, making them unsuitable for studying NM biology or iron–pigment interactions [45]. Although genetically engineered rodent models overproducing NM have emerged, it remains unclear whether their iron levels faithfully reflect the human condition [8, 10, 46]. NM accumulation appears to depend strongly on lifespan and evolutionary proximity to humans, with great apes exhibiting the closest resemblance to human NM [45].

In the present study, we bridge these gaps by quantifying iron and NM concentrations of DN using microscopic colorimetry, μXRF and ultra-high-resolution, ultra-high-field qMRI in ethically collected *post mortem* chimpanzee brains [47, 48]. We systematically quantified iron and NM in large ensembles of DN across the lifespan and linked these microscopic cellular measurements with macroscopic MRI metrics, see Fig. 1. We further connect the NM-iron-saturation across the lifespan to cellular iron toxicity by modeling the equilibrium between different iron binding sites and the labile iron pool.

**Figure 1.**
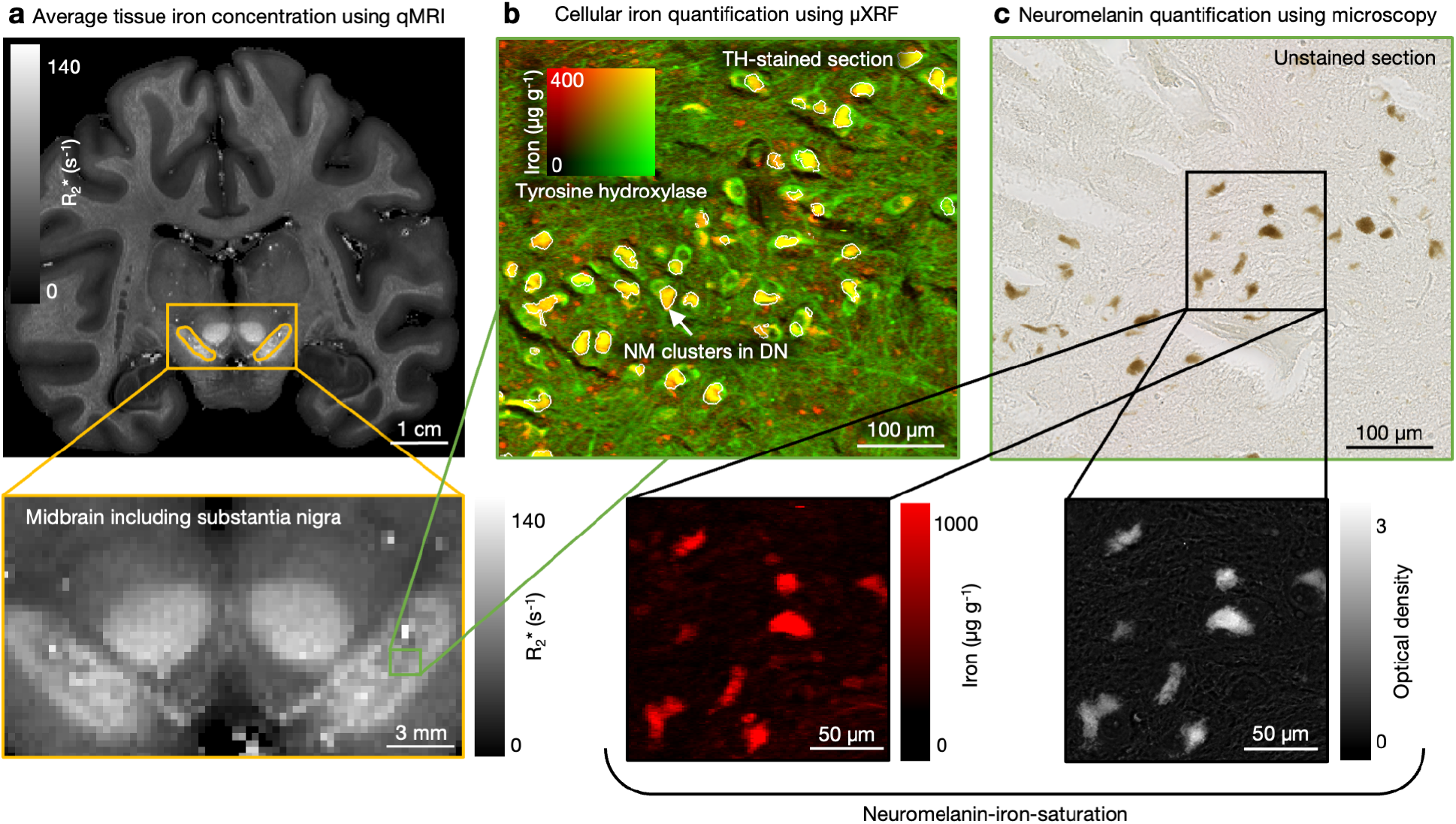
Experimental approach for iron and NM quantification in DN across macro- and micro-scale. **a** Average iron concentration in SN tissue was quantified using iron-sensitive quantitative MRI parameter 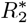 in *post mortem* brains of 20 chimpanzees. The SN can be seen as a hyperintense elongated region on 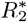-map (yellow delineation, 45-year-old, case S). **b** Cellular iron concentrations were measured in NM of DN in five chimpanzee brains across the lifespan. In the adult brains, iron in DN can be seen as yellow/orange regions in the μXRF iron (red). DN are visualized in nickel-enhanced immunohistochemically-stained sections for tyrosine hydroxylase (TH, green). NM clusters inside the neurons were segmented using a trained classifier on nickel and sulfur maps as NM is known for elevated sulfur concentrations [16, 40](white contour lines, 44-year-old case Q/5). **c** Pigmented DN can be seen in unstained sections as brown regions due to presence of NM (44-year-old case Q/5). NM concentrations were quantified based on optical density maps using the reported extinction coefficients of NM. Cellular iron and NM concentrations of the same neurons were then used to estimate the neuromelanin-iron-saturation.

## 2 Results

The study was conducted in three steps. First, we validated the chimpanzee SN as a suitable model for studies of NM and iron accumulation in human SN. We qualitatively compared NM accumulation stages and NM biology using histology in chimpanzees and humans (section 2.1). We quantitatively compared the qMRI iron marker 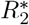 averaged across the SN in both species to characterize the brain iron accumulation (section 2.2). Second, we used this newly established model to understand the potential contribution of iron and NM to age-related neuronal vulnerability. We characterized the lifespan trajectories of iron (section 2.3), NM and the NM-iron-saturation (section 2.4) across large populations of DN using a novel combination of μXRF and optical microscopy. Third, we developed a model linking the NM-iron-ratio of individual cells to toxic labile iron in DN (section 2.5).

### 2.1 Neuromelanin accumulation stages in chimpanzees are similar to those in humans

Qualitative analysis of NM pigmentation in DN of chimpanzees of different ages revealed developmental pigmentation stages similar to those reported in humans [23, 24] (Fig. 2). Three known stages were identified using optical microscopy of unstained sections of substantia nigra *pars compacta* (SNpc), a region characterized by a high density of pigmented DN:

**Figure 2.**
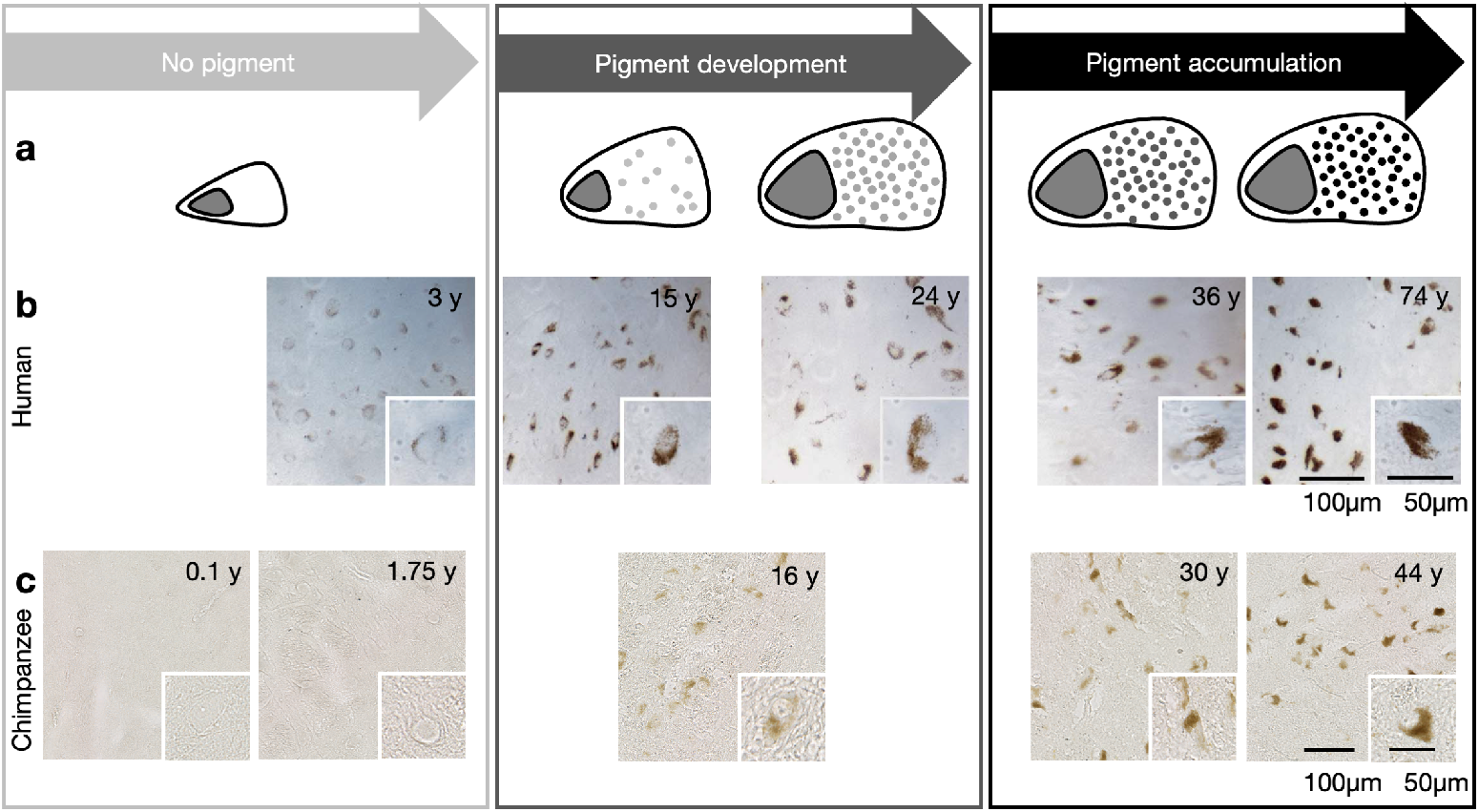
Neuromelanin accumulation in dopaminergic neurons across the lifespan in chimpanzees was similar to humans. **a** Pictograms and **b** optical microscopy of unstained sections of SNpc showing the pigmentation stages previously described in humans (Reprinted from Neurobiology of Aging, **27**, Fedorow et al., “Evidence for specific phases in the development of human neuromelanin”, 506 - 512, Copyright 2006, with permission from Elsevier [24]). **c** The NM pigmentation of DN in chimpanzees (current study), revealed similar stages: no NM until the age of three years, followed by a period of pigment development and a period of pigment accumulation [23, 24].

*Stage 1. No pigment* : The sections of the two youngest chimpanzees (0.1 and 1.75 years) appeared nearly transparent and did not show any pigmentation (Fig. 2c). This observation aligns with findings in humans (Fig. 2b), where NM granules typically begin to appear around three years of age.

*Stage 2. Pigment development* [23]: Light brown areas were observed in the neuronal cytosol of the adolescent chimpanzee brain (16 years, Fig. 2c). This stage falls into the period of NM formation in humans beginning in childhood and continuing until approximately 20 years of age (Fig. 2b).

*Stage 3. Pigment accumulation* [23]: Dense brown NM clusters were present in the adult chimpanzee brains (30 and 44 years, Fig. 2c). These clusters formed goblet-shaped, dark regions partially surrounding the nucleus. The oldest brain (44 years) exhibited the largest and darkest pigmented areas (Suppl. Fig. 11), consistent with human observations, where NM pigmentation intensifies after the age of 20 years (Fig. 2b).

Taken together, the pigmentation and morphology of NM in DN across the chimpanzee lifespan closely correspond to the stages of NM pigmentation previously described in humans, supporting its use as an model for lifespan changes in the human SN. This is different to most other animal models, such as rodents, which lack NM naturally.

### 2.2 Accumulation of nigral iron in chimpanzees is similar to humans

To compare the lifespan accumulation of iron in the entire SN of chimpanzees to humans, we used the iron-sensitive effective relaxation rate 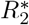. It is approximately linearly related to overall tissue iron content averaged across all cell types and tissue components [41, 49–51]. 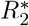 maps can be easily obtained in larger cohorts, enabling a precise characterization of the lifespan changes by dense sampling, although lacking cellular specificity. Ultra-high resolution 7T qMRI (300 μm isotropic) on 20 *post mortem* brains (ages from 0.1 to 56 years) was compared to published ultra-high resolution 7T qMRI (500 μm isotropic) on 105 human participants (18 to 80 years [52]) and published histological iron measures in 52 human donors (0 to 98 years [35]).

The median 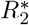 in the chimpanzee SN steadily increased across the lifespan from about 10 s^−1^ at birth to 92 s^−1^ in late adulthood, indicating an accumulation of iron, Fig. 3, Suppl. Fig. 7. The mean 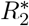 in adult chimpanzee SN (68 ± 15 s^−1^, age *>* 18 years) were close to those in humans (69 ± 8 s^−1^ [52], Fig. 3b) indicating similar levels of adult nigral iron in both species. To characterize the lifespan dynamics of nigral iron, we modeled the age-dependent 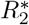 using the exponential saturation suggested for the brain iron accumulation by Hallgren and Sourander [35], and employing Bayesian modeling:

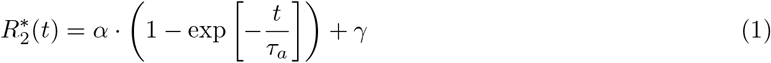

where *t* is the age in years, parameter *α* describes the amplitude of overall changes in 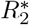 throughout the lifespan, parameter *τ*_*a*_ is the time constant of the exponential saturation, and the intercept parameter *γ* represent the values of 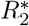 in SN at birth. Note that the function, originally used to model the age-dependence of iron concentration can also be employed to model 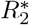, as the linear relation between iron concentration and 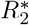 preserves the functional form of age-dependence. Accordingly, the values of parameter *τ*_*a*_, determined for iron dynamics in humans using chemical assays, can be directly compared with our results from fitting the 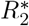 dependence. The maximum of the posterior estimate yielded a time constant *τ*_*a*_ = 38 years (50% highest density interval (HDI): 28 – 47 years, Bayes factor (BF) for *τ*_*a*_ = 38 years: 1.7, Suppl. Fig. 12). The estimated time constant was larger than the time constants reported for human brain regions (10 to 25 years in cortical and subcortical regions as measured histologically [35], and 10 to 13 years in SN as measured with qMRI [53]).

**Figure 3.**
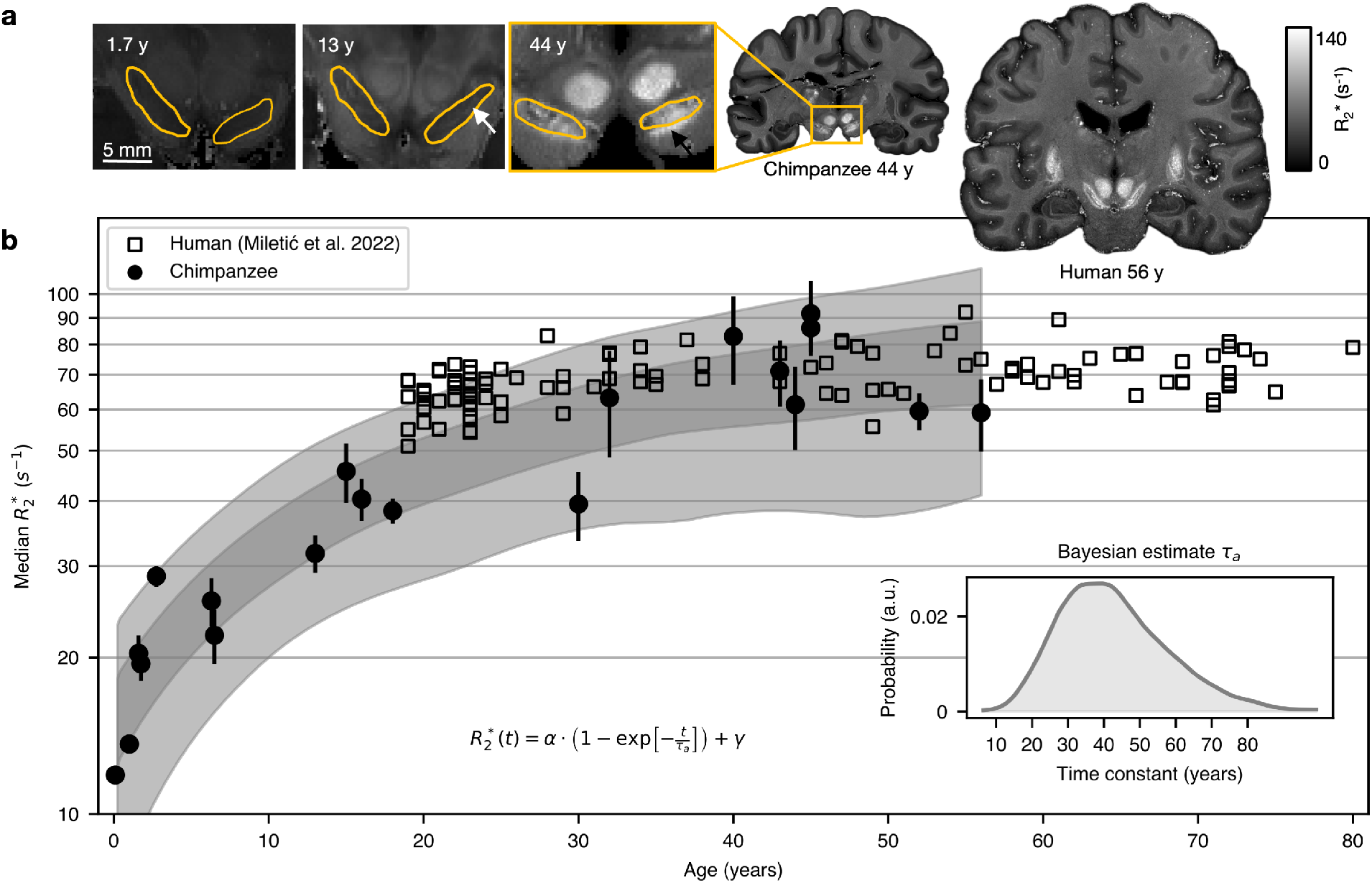
Average tissue iron levels in adult chimpanzee SN were similar to those in human SN as assessed by iron-sensitive MRI. **a** Age-related increase in 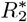 in chimpanzee SN (outline in yellow). 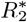 in the SN was low and homogeneous until the age of 13 years. In early adulthood, hyperintense substructures were visible in SN (white arrow), potentially corresponding to the chimpanzee analogues of nigrosomes. From the age of 40 years, high 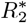 rates in background tissue in SN (black arrow) and in nucleus ruber indicated iron accumulation in these structures. **b** The median 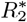 in chimpanzee SN increased across lifespan (circles with SD) and reached a similar level in adulthood as observed in humans (squares [52]). The lifespan trajectory is modeled using the exponential saturation function in a Bayesian approach (50 and 90% highest density intervals (HDI) as gray shaded area) and posterior distribution of the time constant *τ*_*a*_ is shown in the inset.

In addition, variations in the lifespan iron dynamics within the SN were revealed by fine-grained spatial variation in 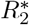, detected by the ultra-high resolution of qMRI (Fig. 3a, Suppl. Fig. 6). While no intra-regional variation was observed before the age of 13 years, localized hyperintensities emerged within the SNpc at older ages, Suppl. Fig. 7, resembling the spatial patterns observed in the human SNpc [54]. This finding suggests that iron accumulates faster in the DN-rich SNpc than the rest of SN.

In summary, the adult iron levels and lifespan dynamics of iron accumulation in the SN of chimpanzees were comparable to those observed in humans. Thus, the chimpanzee represents an informative animal model for studying age-related iron accumulation in the human SN.

### 2.3 Dopaminergic neurons are particularly fast in accumulating iron across the lifespan

We developed a high-throughput pipeline for cellular iron quantification in a large group of neurons by combining μXRF with advanced histology and machine-learning–based image analysis, Fig. 1, Suppl. Fig. 8. Tissue sections from SNpc were stained using nickel-enhanced TH-immunohistochemistry to identify DN [36, 55]. The high-brilliance synchrotron X-ray source PETRA III at the Deutsches Elektronen-Synchrotron DESY enabled quantitative mapping of iron, nickel, and sulfur concentrations with microscopic resolution. This resolution was sufficient to segment individual cells while covering several square millimeters of tissue, Fig. 4, Suppl. Fig. 8, 9. This approach enabled the analysis of large neuronal populations, thereby bridging the gap between cellular-level iron quantification and macroscopic MRI-based iron metrics. DN somata were automatically segmented in nickel maps. Conversely, NM clusters were segmented in sulfur maps, as NM is known for elevated sulfur concentration due to its benzothiazine rings and cysteine-containing residues (Suppl. Fig. 9) [16, 40]. The NM-segmentation was restricted to the DN-somata mask to quantify the mean iron concentrations within NM clusters of DN.

**Figure 4.**
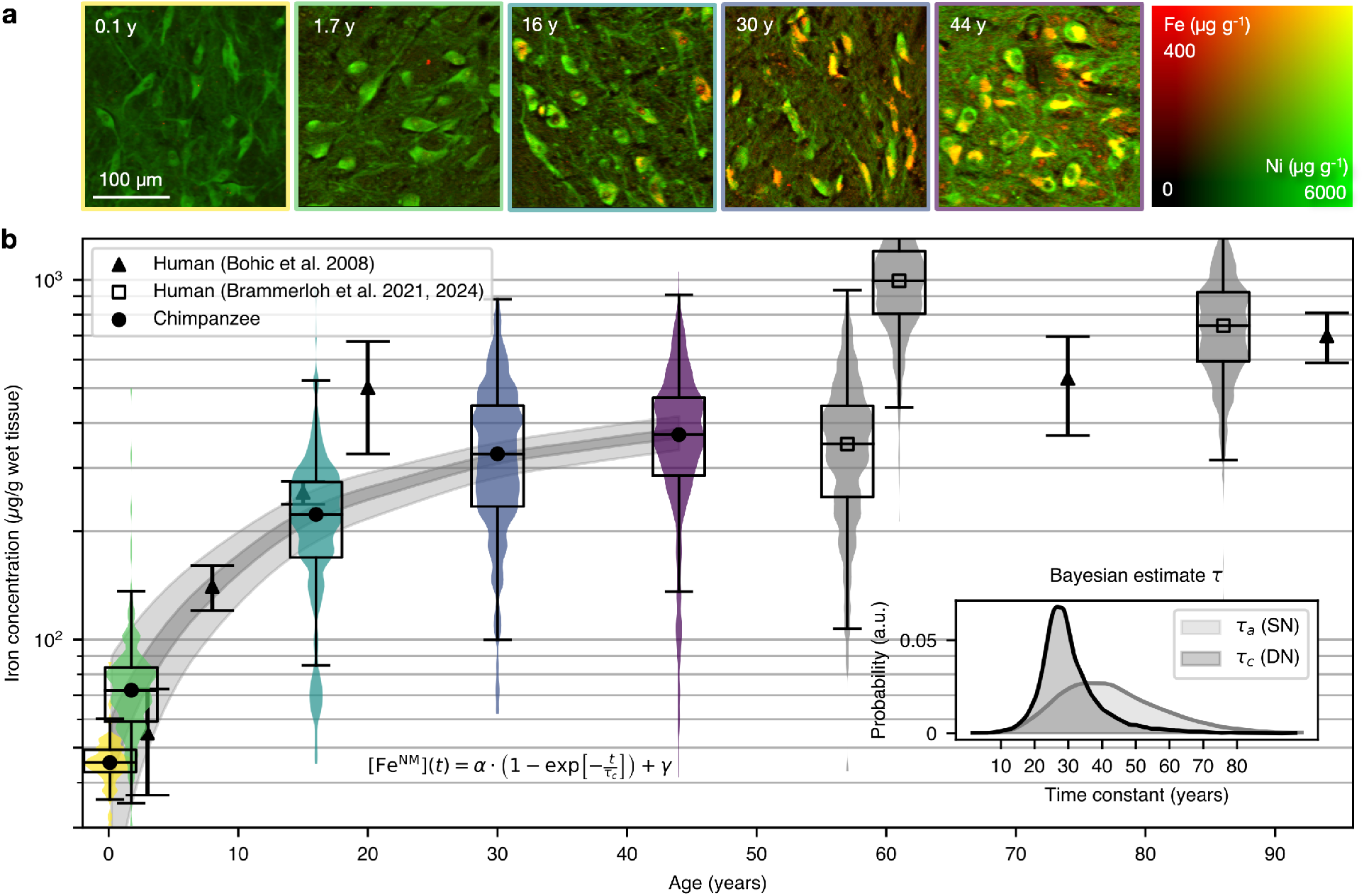
Iron concentration in dopaminergic neurons show faster lifespan accumulation than the average iron in the substantia nigra. **a** Quantitative iron (red) and nickel (green) maps of nickel-enhanced immunohistochemically stained DN. Iron accumulation in NM in DN is visible as yellow/orange clusters in older animals. **b** The age-dependent iron accumulation in NM in DN in chimpanzees (filled circles, violins) and modeling results across the lifespan are shown. Sparse reports of iron concentrations in human DN are shown for comparison (squares, gray violin plots [41, 44], triangles with error bars [40]), demonstrating similar values to those in chimpanzees. The iron concentrations in [40] were multiplied by a factor of 0.9 to correct for the tissue matrix and convert to units of iron concentrations to μg g^−1^ wet tissue weight (based on personal communication with the authors). The violin and box plots illustrate the distribution of mean iron concentrations measured in individual cells, their median (symbol and horizontal line in the boxes), and interquartile ranges. The resulting 50% and 90% highest density intervals are shown as gray shaded areas. The inset shows the posterior probability of the time constant for the cellular iron concentrations *τ*_*c*_ in comparison to the one for the average tissue iron *τ*_*a*_ from Fig. 3.

In a subset of five animals covering the entire chimpanzee lifespan (ages 0.1 to 44 years, Table 3), we measured the mean iron concentration in a set comprising 91 to 698 automatically segmented NM clusters per animal, Table 1, Suppl. Fig. 9. The iron concentration in NM clusters increased eightfold from birth to adulthood, Fig. 4. Iron concentration in tissue outside of the DN also increased across the lifespan, albeit to a lesser extent, raising by a factor of approximately two from birth to adulthood, Table 1. Potential biases introduced by sample preparation and segmentation were quantified and found to be small relative to the biological variability of iron concentration within and between individuals, Suppl. Fig. 10.

We modeled the age-dependent cellular iron concentration in NM clusters [Fe^NM^] as a function of age *t* by the exponential saturation model using chimpanzee data and a Bayesian approach [35]:

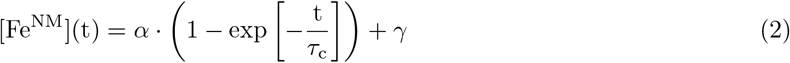

The results of the Bayesian analysis and the posterior distributions are shown in Fig. 4 for the time constant parameter *τ*_*c*_ and in Suppl. Fig. 12b for other model parameters. The maximum posterior probability for the time constant *τ*_*c*_ was 27 years (50% HDI: 23 – 31 years, Bayes factor 3.4). This value was significantly shorter than the time constant of 38 years determined for the lifespan dynamic of averaged iron concentration in SN (Fig. 4 inset), as confirmed by Bayesian analysis of posterior probability distributions, see Suppl. Fig. 12c.

This provides evidence that iron in the pigmented DN accumulates faster than the iron across other cellular and extra-cellular tissue components of the SN. Consequently, these neurons not only reach higher cellular iron levels but are also exposed to elevated iron concentrations earlier in life than other cell types.

### 2.4 Neuromelanin-iron saturation remains nearly constant across lifespan and between species

We developed a novel method to quantify NM concentrations and the NM-iron-saturation (the saturation of NM with iron) in individual dopaminergic neurons by combining colorimetric analysis of optical microscopy with single-cell iron quantification. NM concentration of each DN was derived from optical density (OD) maps calibrated using the known extinction coefficients of pheomelanin and eumelanin, the main chromophores of NM, together with their established mass fractions within the NM pigment, see section 4.7 [7, 56, 57]. A linear relationship between OD and sulfur concentrations supported the assumption that the OD is modulated by NM (Suppl. Fig. 11a).

Cells were segmented using a trained classifier on the optical microscopy images, which were co-registered to the corresponding iron maps (Fig. 5). We applied this method to brain tissue from three older chimpanzees (16, 30 and 44 years) exhibiting detectable NM pigment, as well as to published data from two human donors (61 and 86 years [41, 44]).

**Figure 5.**
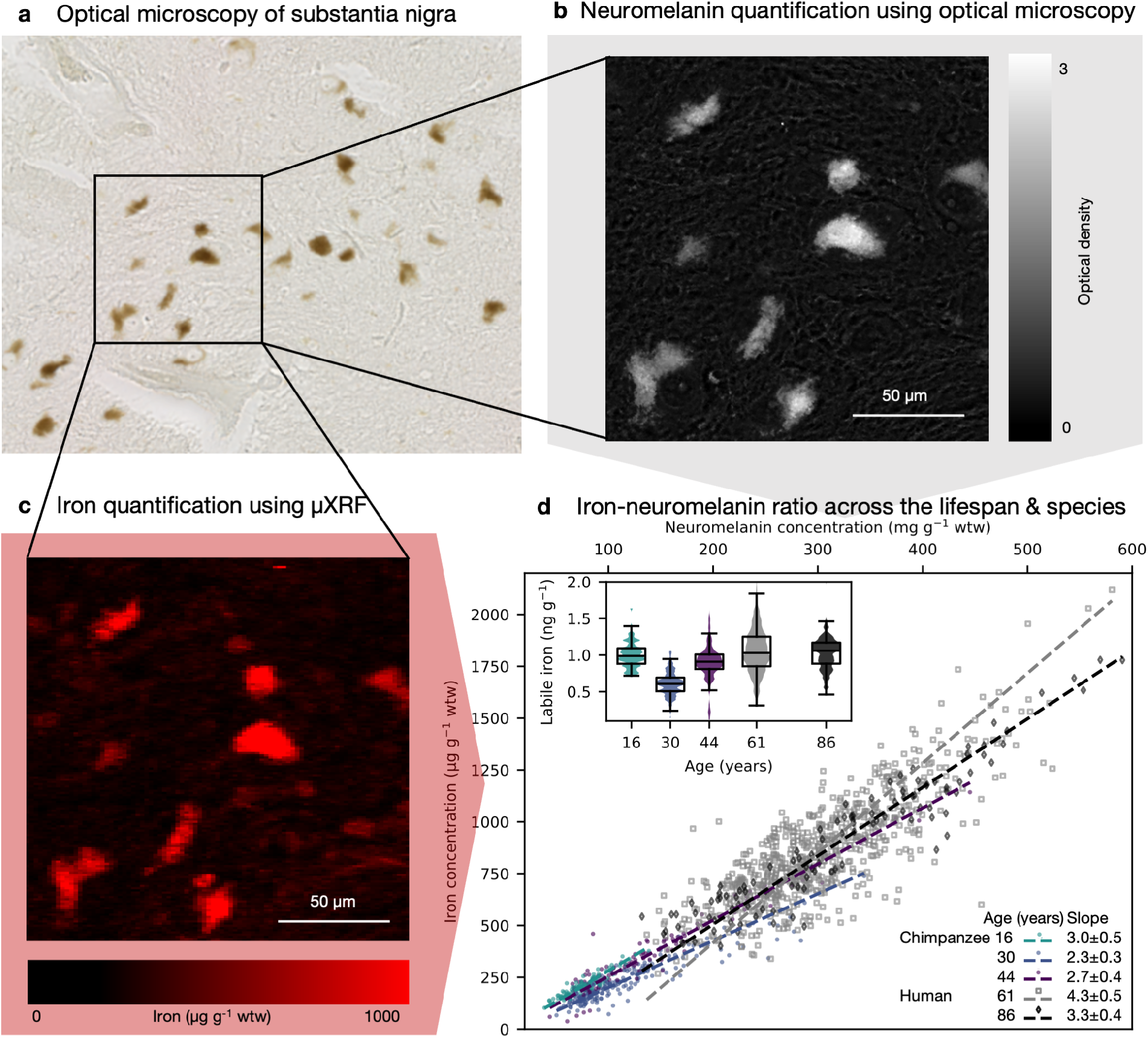
The NM-iron-saturation estimated on the cellular level remained stable across the lifespan and across species. **a:** Pigmented dopaminergic neurons can be seen as brown regions in optical microscopy of the substantia nigra (44-year-old). **b:** NM concentration was quantified using optical density and the reported absorption coefficients of pheomelanin and eumelanin [57]. **c:** The quantitative iron map of the 44 year-old chimpanzee shows elevated iron concentrations in the pigmented neurons. **d:** We observed NM-iron-saturation of about 2.3 to 4.3 μg iron per mg NM for chimpanzees and humans. The human samples from previous studies were partly reanalyzed [41, 44]. The inset shows the estimation of labile iron concentrations which remains stable across the lifespan.

We observed a significant increase in median NM concentration with age rising from 74 ± 18 mg g^−1^ in the adolescent to 96 ± 53 mg g^−1^ and 126 ± 77 mg g^−1^ in the adult chimpanzee brain, Suppl. Fig. 11b (ANOVA NM(age): df = 2, F = 26, p *<* 0.001) and even significantly higher NM levels in the elderly humans (292 ± 71 mg g^−1^ and 309 ± 103 mg g^−1^, ANOVA NM(species): df = 1, F = 1740, p *<* 0.001). The iron and NM concentrations in individual DN showed a strong linear relationship in all individuals (by ascending age: Pearson’s r = 0.923, 0.925, 0.972, 0.820, 0.951.

The slope of the orthogonal distance regression of this intra-individual linear relationship was used to determine the NM-iron-saturation for each individual. While iron concentrations increased almost fourfold between the youngest chimpanzee and oldest human, the NM-iron-saturation varied by only 40% across ages and between the species, ranging from 2.3 to 3.0 μg iron mg^−1^ NM in chimpanzees and from 3.3 to 4.3 μg iron mg^−1^ NM in humans.

A generalized linear model describing the single-cell iron concentration as a function of NM, age and species – including NM *×* age and NM *×* species interaction terms – showed a significant main effect of NM (p *<* 0.001) and significant interaction between NM and species (p = 0.001), whereas the interaction between NM and age was not significant (p = 0.155). These results indicate a 40% higher NM-iron-saturation in humans compared to chimpanzees, with no significant increase of NM saturation with age. This conclusion was supported by Bayesian ANCOVA (section 4.7). However, our dataset is sparse and not balanced across species due to younger individuals in chimpanzees and the absence of young adult human data. Broader age sampling and overlapping age ranges between the two species are required to estimate age-related and inter-species effects with higher precision. Nevertheless, the NM–iron-saturation varied far less across the lifespan than the absolute levels of NM and iron, indicating efficient iron chelation across the lifespan with no evidence of NM oversaturation.

### 2.5 Modeling iron-neuromelanin toxicity in dopaminergic neurons

Finally, we linked the measured NM-iron-saturation to intracellular iron toxicity by modeling the chemical equilibrium between NM-bound iron and the cytosolic labile iron pool. Assuming reversible exchange and equilibrium between cytosolic iron and the two iron binding sites within NM – the high-affinity site (dissociation constant *K*_*L*_ = 94.31 nM) and the low affinity site (*K*_*H*_ = 7.18 nM) [22] – we modeled the concentration of labile iron as a function of total cellular iron and NM concentration. We showed analytically, that the concentration of labile iron depends solely on the total iron bound in NM divided by its maximum iron-binding capacity of NM, 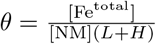 . *L* and *H* are maximum possible mass of bound iron at low- and high-affinity bindings sites in NM per mass unit of NM respectively (*B*_*max*_ in [22]). The cellular concentration of labile iron is determined by:

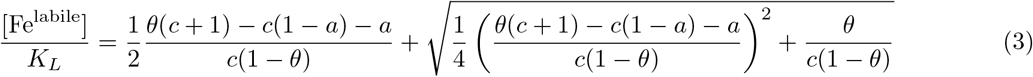

where *a* and *c* are constants describing chemical properties of iron binding sites of NM, with 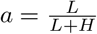 being the mass fraction of iron in low-affinity binding site, in fully saturated NM and 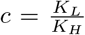 being the ratio of dissociation constants of iron in low- and high-affinity binding sites. Thus, the empirically estimated NM–iron saturation ratio provides a direct quantitative link to intracellular iron toxicity, if all model parameters in Eq. 3 are known.

In titration experiments with extracts of human NM the model parameters *a* and *c* were estimated to be *a* ≈ 0.84 and *c* ≈ 13 [22]. *θ* was estimated using a novel technique in this study for each individual cell. The only unknown parameter is the total amount of iron in fully saturated NM, [Fe^max^] = [NM](*L* + *H*), as literature estimations of these parameters are controversial, varying between 1.1 μg and 11.3 μg per mg NM in different studies [22, 38].

Using the proposed model, the literature values for NM-iron-binding properties and assuming an iron-binding capacity of 11.14 μg iron per mg NM (factor 10 from the maximally bound iron in [22], in agreement with other extracted NM studies [38]), we estimated the concentration of labile iron for three chimpanzee and two humans (Fig. 5d). Very similar, low levels of labile iron were detected in all individuals, pointing towards efficient iron chelation. With the exception of the 30-year-old chimpanzee, which showed lower labile iron concentrations, the median of labile iron varied less than 10% across individuals and species.

For the first time, we provide a quantitative basis and a theoretical framework at the cellular level to estimate the labile iron concentration within individual DN.

## 3 Discussion

In this study, we investigated how the accumulation of iron and NM contributes to the selective vulnerability of DN in the SN. Using a novel combination of ultra-high resolution quantitative MRI, μXRF, optical imaging, and automated segmentation, we characterized the lifespan trajectory of cellular iron and NM concentrations in *post mortem* brains of chimpanzees and humans (Fig. 1). Our approach allowed the analysis of an unprecedented number of individual neurons, yielding representative and specific results of cellular iron concentrations. By modeling the iron binding in NM, we identified NM-iron-saturation as the key parameter for estimating the levels of toxic labile iron.

The NM-iron-saturation remained remarkably consistent across the lifespan in healthy individuals. In other words, NM accumulation keeps pace with increasing cellular iron levels, thereby maintaining its neuro-protective function and preventing saturation. In line with a neuroprotective effect, our chemical equilibrium model shows that the labile iron concentrations in individual cells is constant for constant NM-iron-saturation. Our findings and developed methods are of importance because previous studies only examined NM-iron-saturation in extracted NM, where the iron load is altered through the extraction procedure and then pooled across many cells and several individuals [22, 37, 38, 58]. Our results help to resolve the controversy on mechanisms of neuronal vulnerability in context of PD by demonstrating that NM typically remains unsaturated [8] rather than becoming iron overloaded [16, 59].

These insights relied on a unique resource: a lifespan collection of ethically sourced *post mortem* chimpanzee brains. This collection overcomes a major limitation in human *post mortem* research: the ethical and practical difficulty of obtaining brain tissue from children and young adults. We demonstrated that the chimpanzee is a more informative model for studying the lifespan trajectory of DN in the human SN than traditional laboratory animals, which do not accumulate NM naturally, have low brain iron levels, and short lifespans [10, 45, 46]. In contrast, chimpanzees have a lifespan comparable to humans [60]. Moreover, they show similar stages of NM accumulation (Fig. 2) and comparable tissue and cellular iron levels in the adult SN (Fig. 3, 4).

We developed and applied a novel combination of advanced biophysical methods to bridge the gap between cellular- and tissue-level iron trajectories. Benefiting from recent advances in resolution and brilliance of synchrotron μXRF, we performed quantitative cellular iron mapping in macroscopically large areas. Automated segmentation of many DN in these maps using a trained classifier enabled the characterization of the biological variability of cellular iron concentrations in the SN. Ultra-high field and ultra-high-resolution quantitative MRI allowed the quantification of average tissue iron in the SN across a larger group of animals. This facilitated a comparison of the lifespan trajectory of cellular iron to average tissue iron and bridging between cell-specific histological measures and non-invasive MRI measures. As the labile iron pool can be barely measured directly [61], we estimated its concentration by modeling the iron binding within NM and the NM-iron-saturation, Fig. 5.

Overall, the iron concentrations of NM clusters in DN in the chimpanzee SN obtained in our study (Table 1) matched previously reported data in humans (Table 2). Both our cellular (46 to 371 μg g^−1^) and extraneuronal iron concentrations (35 to 80 μg g^−1^) fell well within the range of values reported in the literature for human tissue (55 μg g^−1^ to 805 μg g^−1^ for cellular and 21 to 200 μg g^−1^ for extraneuronal iron, Table 2). Note that previous studies encompassed both frozen and fixed tissue samples, a factor which may partially explain the observed variations in iron distribution.

**Table 1.** Iron concentrations in individual NM clusters (median and standard deviation of mean iron concentration in individual NM clusters) measured with μXRF for five chimpanzee of different ages. The iron concentration in tissue outside of NM (extraneuronal iron) and the number of individual NM clusters is shown. The median iron concentration in NM clusters (μg g^−1^ wet tissue weight (wtw)) increased with age by a factor of eight, while the iron concentration in the surrounding tissue only increased by factor two. Measurements were performed across macroscopic neuron-dense regions of the SNpc.

| Case | 1 | 2 | 3 | 4 | 5 |
| --- | --- | --- | --- | --- | --- |
| Age (years) | 0.1 | 1.75 | 16 | 30 | 44 |
| Iron concentration (median ± SD, $\mu\text{g/g}$) | $46 \pm 7$ | $72 \pm 38$ | $223 \pm 126$ | $328 \pm 156$ | $371 \pm 159$ |
| Number of segmented NM clusters | 91 | 270 | 159 | 303 | 698 |
| Extraneuronal iron concentration (median ± SD, $\mu\text{g/g}$) | $34 \pm 36$ | $45 \pm 26$ | $42 \pm 37$ | $57 \pm 35$ | $79 \pm 51$ |

**Table 2.** Summary of studies reporting iron concentrations in the human SN and individual DN (μg g^−1^ wet tissue weight). N: number of individuals; AAS: atomic absorption spectrophotometry; NAA: neutron activation analysis; μXRF: X-ray fluorescence; PIXE: proton-induced X-ray emission; ICP-MS: inductively coupled plasma mass spectrometry. ^*a*^ Iron concentration in dry tissue weight divided by 5. ^*b*^ Iron concentration multiplied by factor 0.9 to correct for tissue matrix.

| Study | [Fe] ($\mu\text{g g}^{-1}$) | | | Age range (y) | N | Method | Tissue | Ref |
| --- | --- | --- | --- | --- | --- | --- | --- | --- |
|  | in SN | in DN | outside DN |  |  |  |  |  |
| Lifespan | 185 ± 65 | – | – | 30 – 100 | 52 | Colorimetry | unfixed | 35 |
|  | 40 – 160 | – | – | 0 – 80 | 33 | AAS | freeze-dried | 62<sup>a</sup> |
|  | 21 – 199 | – | – | 1 – 90 | 18 | NAA | – | 12 |
| | – | 55 – 700 | 40 – 200 | 3 – 94 | 6 | $\mu\text{XRF}$ | fixed | 40<sup>b</sup> |
| Cellular iron | – | 100 | 60 | – | – | $\mu\text{XRF}$ | freeze-dried | 42 |
| | 140 | 360 | – | 82 ± 10 | 8 | PIXE | fixed | 43<sup>SI<sup>a</sup></sup> |
| | – | 221 | – | 82 ± 6 | 5 | PIXE | frozen | 43<sup>SI<sup>a</sup></sup> |
| | – | 427 | – | 82 ± 10 | 10 | $\mu\text{XRF}$ | fixed | 43<sup>SI<sup>a</sup></sup> |
| | – | 253 | – | 82 ± 6 | 5 | $\mu\text{XRF}$ | frozen | 43<sup>SI<sup>a</sup></sup> |
| | – | 490 | 200 | 66 ± 16 | 8 | PIXE | fixed | 36 |
|  | – | 805 | 230 | 68 | 3 | PIXE | fixed | 41 |
| Bulk iron | 146 | – | – | 89 ± 4 | 2 | AAS | freeze-dried | 63 |
|  | 50 | – | – | 81 | 8 | ICP-MS | – | 64 |
| | 200 ± 50 | – | – | 78 | 1 | $\mu\text{XRF}$ | – | 32 |
| | 99 ± 48 | – | – | 74 | 18 | ICP-MS | dried | 65<sup>a</sup> |

By analyzing 91 to 698 dopaminergic neurons per case (Fig. 4) – far more than the 5 to 30 cells sampled in earlier studies [40] – we showed that the high variability of previously reported neuronal iron concentrations likely reflects the underlying broad biological distribution. Here, we provide a more representative sampling of cellular iron distribution across large cell ensembles.

To compare the lifespan trajectory of cellular iron concentration measured by μXRF with bulk tissue iron concentrations, we employed quantitative *R*_2_^*∗*^ maps with sub-millimeter resolution across a larger group of *post mortem* brains, see Fig. 3. We found that the iron accumulation in the chimpanzee SN progressed significantly slower than in individual DN, as indicated by a longer time constant (*τ*_*a*_ = 38 years in the SN as compared to *τ*_*c*_ = 27 years in DN). This difference underscores the distinct mechanisms of iron regulation within DN compared to other cells, resulting in higher iron levels and accelerated accumulation over the lifespan. Consequently, average 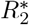 values in the SN cannot serve as a reliable proxy for the specific iron concentration in DN.

The obtained time constant of iron accumulation in the chimpanzee SN was larger than the MRI-based constant of 13 ± 6 years for the time constant of average 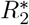 in the human SN [53] and the range of time constants of 10 to 25 years reported across subcortical regions based on histology in human *post mortem* tissue [35]. This may point to an overall faster iron accumulation in humans compared to chimpanzees.

To answer the question of whether the drastic increase in cellular iron in aging neurons is linked to increasing iron toxicity, we analyzed the NM-iron-saturation and estimated the concentration of potentially toxic, labile iron. The obtained values for NM-iron-saturation were in the order of magnitude of those reported in the literature. The NM-iron-saturation in chimpanzee and human SN (mean 3.1 ± 0.7 μg iron per mg NM) were slightly lower than the reported averaged NM-iron-saturation in extracted humans NM, which range from 5.2 [22] to 9.2 [58] and 11.3 μg iron per mg NM [38]. Because we found lower NM-iron-saturations than previously reported, the NM clusters examined in this study appear unsaturated, hence capable of further iron chelation. Higher NM-iron-saturation values in extracted NM, as compared to our values *in situ*, could be explained by an iron uptake during the isolation process of NM [37, 38].

Despite the eightfold increase in iron concentration in DN from birth to adulthood, the NM-iron-saturation remained similar throughout the adult lifespan. As a consequence, the labile iron concentration likewise remained at a similarly low level, at about 1 ng g^−1^ throughout lifespan, Fig. 5d. Only one study estimated labile iron concentrations in tissue of healthy controls and PD patients in homogenized samples of SN tissue, reporting higher values, as compared to our findings: 90 ± 9 ng g^−1^ in PD, vs. 37 ± 2 ng g^−1^ in control [18]. Other authors have doubted that labile iron can be measured reliably in tissue homogenates [61]. Using our novel approach, we estimated the cellular labile iron concentrations without extraction, potentially providing more biologically plausible and unbiased values. Our quantitative experimental approach can be translated to examine changes in the iron binding of NM in disease [15].

The methods developed in this study offer novel applications beyond the scope of iron-NM dynamics. For instance, by demonstrating a linear relationship of NM concentrations and iron concentrations in DN, our data supports the method for estimating the cellular iron concentrations in NM in DN using only optical microscopy used in a previous study [41]. Moreover, the cellular iron trajectories established across the lifespan can directly inform the development of next-generation MRI biomarkers for DN integrity. So far, biophysical models linking microscopic iron concentrations to macroscopic MRI signal have been limited to data from elderly patients and could now be extended using our dataset covering the entire lifespan.

This work is subject to several limitations, some of which are intrinsic to *post mortem* research. First, while the iron concentration in chimpanzee SN was measured in 20 individuals, cellular measurements were limited to only five individuals due to limited synchrotron beamtime. To make full use of these sparse data, we used Bayesian modeling with literature-informed priors. Second, we used 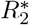 as a proxy of tissue iron, while it is also known to be sensitive to myelin. However 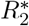 remains a robust proxy for average iron in myelin-poor brain regions like the SN [30]. When comparing *post mortem* to *in vivo* MRI, note that formalin fixation may increase the effective relaxation rate of tissue by 14% [66]. Nonetheless, the fixation process was standardized for all samples, hence we expect a similar change in relaxation rates for all chimpanzee brains. Third, the spectroscopic estimation of NM concentration relies on literature absorption curves of eumelanin and pheomelanin, and reports on human NM composition [56, 57]. These parameters may vary across species and throughout the lifespan, potentially introducing biases into the estimates of NM. Finally, our chemical equilibrium model utilized iron-binding constants derived from a single study [22]. Future replications may provide updated constants to refine our model estimations. The assumption of a chemical equilibrium between NM-bound iron and the labile iron pool is supported by recent chelation trials demonstrating that iron can be successfully sequestered [25, 26]. Our model assumes that most of the iron in NM clusters is bound to NM, neglecting transferrin- and ferritin-bound iron. This is justified by our observation that iron concentrations in the surrounding tissue – likely reflecting transferrin- and ferritin-bound iron – were significantly lower than those within the DN. Additionally, NM is reported to be encapsulated within lipid membranes and isolated from the cytosol, preventing direct contact with transferrin and ferritin.

Combining the cellular specificity of μXRF with non-invasive qMRI, we have bridged the micro- and macroscale trajectories of lifespan changes in the SN. Introducing the chimpanzee SN as an informative model of iron accumulation in the human SN, we demonstrated that while DN accumulated iron significantly faster than surrounding tissue, this iron remained effectively chelated by NM throughout the healthy lifespan. Consequently, labile iron concentrations remained remarkably low. These findings provide a much-needed reference for the lifespan trajectory of NM-iron-saturation and the intracellular labile iron concentrations, contributing to the understanding of vulnerability of DN to neurodegeneration with age. Moreover, the methods and findings presented in this study establish a quantitative framework that can inform the development, targeting, and mechanistic evaluation of iron-chelating therapies in Parkinson’s disease.

## 4 Materials and methods

### 4.1 Tissue samples

The 20 *post mortem* whole brains of chimpanzees (Pan troglodytes) who died of natural causes were collected in African field sites, sanctuaries and European zoos using an ethical pipeline [47] within the multidisciplinary project *Evolution of Brain Connectivity*. Detailed information on age, sex, subspecies, cause of death and *post mortem* interval before fixation for all cases is presented in the Suppl. Table 5. The tissue collection protocol adhered to the ethical guidelines established for primatological research at the Max Planck Institute for Evolutionary Anthropology, Leipzig and received approval from the ethics committee of the Max Planck Society. The brains were extracted within 24 h after death and immersion-fixed in 4 % formaldehyde solution prepared in phosphate-buffered saline (PBS) for a period of 8-12 weeks. Fixed brains were transferred to PBS containing 0.1 % sodium azide at pH 7.4 for long term storage, at least 10 days prior to MRI scanning. For a subset of five brain samples (Table 3), the left half of the midbrain was separated and processed for histology and μXRF iron quantification.

**Table 3.** Tissue information for the *post mortem* brain samples from 5 chimpanzees (*Pan troglodytes*). The *post mortem* interval (PMI) denotes the time span between death and brain fixation. Bcbva: Bacillus cereus biovar anthracis.

| Case | 1 | 2 | 3 | 4 | 5 |
| --- | --- | --- | --- | --- | --- |
| Subspecies | schweinfurthii | verus | verus | schweinfurthii | mixed |
| Age (years) | 0.1 | 1.75 | 16 | 30±5 | 44 |
| Sex | m | m | m | m | f |
| Brain weight (g) | 129 | 340 | 347 | 361 | 357 |
| PMI (h) | 4 | 24 | <16 | 12 | 1 |
| Cause of death | con-specific aggression | Bcbva infection | Bcbva infection | inter-specific aggression | cardiovascular disease |

### 4.2 Quantitative MRI

Whole brain quantitative multi-parametric maps with 300 μm isotropic resolution were acquired on a Siemens Terra 7T whole-body MRI scanner (Siemens Healthineers, Erlangen) using a standardized protocol and a 32 channel receive - 1 channel transmit human head radio-frequency coil (Nova Medical, Wilmington, MA). The brains were embedded inside an acrylic ovoid-shaped container and immersed in the proton-free fluid Fomblin (Solvay Solexis, Bollate, Italy).

Three 3D multi-echo fast low angle shot (FLASH) scans with longitudinal relaxation rate (T1), proton-density (PD) and magnetization-transfer (MT) weighting were acquired with the following parameters, as described in [67]: field of view (FOV) 118.125 *×* 135*×*86.4 mm^3^; matrix size 378 *×* 432 *×* 288 (300 μm isotropic resolution); bandwidth BW = 331 Hz/pixel; twelve equidistant echo times TE between 3.63 ms and 41.69 ms; Δ TE = 3.56 ms; repetition time TR= 70 ms; excitation flip angles = 84^*°*^ (T1-weighted), 18^*°*^ (PD-& MT-weighted); MT pulse characteristics: Gaussian 3 kHz offset, flip angle 700^*°*^. The acquisition of each of the weighted images took about 2 hours. The temperature was kept at 30 ± 2 °C during the scans using warm air flow. It was measured throughout the scans using a fiber optic temperature sensor (LumaSense) using the TrueTemp software (Luxtron).

The static magnetic field *B*_0_ was mapped using a dual gradient echo sequence (matrix size: 96 *×* 96 *×* 64; TR = 1020 ms; TE = 10 and 11.02 ms, flip angle = 20^*°*^). The 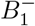 field was corrected using images acquired with the receive coil and the transmit coil in receive mode as described in [67]. The RF transmit field 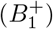 was mapped at an isotropic resolution of 4 mm (matrix size: 64 *×* 48 *×* 48; TR = 500 ms; TE = 40.54 ms; mixing time = 34.91 ms; flip angles 330^*°*^ to 120^*°*^ in steps of 15^*°*^; RF duration 24 μs; GRAPPA acceleration factor 2*×*2).

Quantitative maps of longitudinal relaxation rate R1, effective transverse relaxation rate 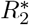, proton density maps, and magnetization transfer saturation maps were calculated using the hMRI-toolbox [67–69]. Quantitative 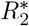 maps of the whole brain were calculated across all three weighted acquisitions by a weighted least square fit implemented in the hMRI-toolbox. The SN was segmented as a myelin-poor region, appearing as a hypointense region in magnetization transfer saturation maps next to the nucleus ruber.

### 4.3 Histology and immunohistochemistry

For the iron quantification experiments using X-ray fluorescence and histology, small tissue blocks of mid-brains, containing the SN with total volumes of approximately (30 mm)^3^ were embedded in paraffin (Histowax, SAV LP, Flintsbach) and cut into 10 μm thin coronal sections using a sliding microtome (Jung Histoslide 2000, Leica, Wetzlar). Sections containing SN pars compacta were mounted on Superfrost^®^Plus glass slides (Thermo Fischer Scientific, Massachusetts, USA) and deparaffinized in xylene.

For each animal, consecutive sections were used for μXRF and two sets of samples were prepared: 1) unstained sections, 2) sections immunohistochemically stained for tyrosine-hydroxylase (TH, rabbit anti-tyrosin hydroxylase antibody AB 152, Merck Millipore, Darmstadt). The regulatory protein tyrosine hydroxylase (TH) constitutes the bottleneck for the dopamine production [11]. Immunohistochemical anti-TH staining labeled cellular structures involved in dopamine synthesis and therefore serves as a reliable marker for DNs. Nickel enhancement with 3,3’-diaminobenzidine (DAB) staining using ultra-pure nickel was performed to make DNs directly visible in μXRF element maps [36, 55]. Transmission light microscopy was collected on all sections with 10*×*, 40*×* or 63*×* Plan-Apochromat objectives using an automated slide scanner microscope (AxioScan Z1 microscope, Zeiss, Jena).

### 4.4 Synchrotron microprobe X-ray fluorescence imaging

The cellular iron concentrations in DNs were quantified using μXRF on the histological sections of five chimpanzees’ midbrains (Table 3) at PETRA III hard X-ray beamline P06 at Deutsches Elektronen-Synchrotron DESY, Hamburg [70]. The sections were embedded in a mounting medium (DePeX, Merck Millipore, Darmstadt), detached from the glass slides and mounted into aluminum frames. Photons with energy of 12 keV were used to ensure the excitation of iron’s inner electrons (photoelectric effect). The photon flux was 3 *×* 10^11^ photons*/*s. The beam size at the sample location was adjusted to 2*×*0.7 μm^2^ with the help of Kirk-patrick–Baez mirrors and prefocusing compound refractive lenses. An energy dispersive silicon drift detector (Vortex ME4, 350 μm Si, SII Nanotechnology, Northridge, USA) with a Beryllium window (Be 12.5 μm) was positioned at a distance of 22 mm (air path) from the sample. The X-rays emitted from the sample onto the detector formed an angle of 225^*°*^ with the direction of the beam.

For each section, a large overview region of interest (ROI) of about 7*×*8 mm^2^ (matrix size 700 *×* 800, 10*×*10 μm^2^ resolution) and two smaller ROIs within SN pars compacta (1*×*1 mm^2^, matrix size 500 *×* 1000, 2*×*1 μm^2^ resolution) were scanned, covering several hundred cells per animal in total. For each case one unstained and one TH-stained section were imaged. For case 5, two additional consecutive sections were imaged. For adult animals (cases 3, 4, and 5) the dwell time was 3 ms. Longer dwell times of 6 ms for case 2 and 12 ms for case 1 were used to achieve high SNR, despite low iron concentration in young brains. Quantitative iron area mass density maps were obtained using a fitting algorithm implemented in PyMca [71] and calibration by comparison with the XRF spectra recorded from a thin-film μXRF reference sample (four-element standard containing Cu, Fe, Cr and Ti on a 200 nm thin Si_3_N_4_ window; type RG-200-0510-C00-X, AXO Dresden GmbH, Dresden). Semi-quantitative sulfur and nickel maps were calculated using the software xraylib [72]. An iron area mass density of 1 ng*/*cm^2^ in the tissue of 10 μm thickness and with a tissue mass density of 1 g*/*cm^3^ corresponds to an iron concentration of 1 μg g^−1^. Note that we have not corrected the element concentrations for volume shrinkage due to paraffin embedding and histological processing, estimated to be 0.76 in a previous study [41].

The iron maps of the 61-year-old human sample (sample 3 in [41]) were acquired using proton-induced X-ray emission (PIXE) as described in [41], while the iron map of the 86-year-old human sample (sample 2 in [41]) was measured in the same μXRF session using the same settings as described for the chimpanzee samples above.

### 4.5 Neuron segmentation using trainable classifier

We developed a pipeline for automated segmentation of the DN and NM clusters within DN, aiming for comparable segmentation performance across individuals of different ages. Such a unified automated pipeline is important, as the size and NM content of neurons and NM clusters vary across developmental stages [23, 24].

Two independent segmentations utilizing the sulfur and the nickel μXRF maps were performed using a trainable random forest classifier (ilastik, v 1.4.0, [73]): (i) segmentation of the neuronal somata (cell bodies) and (ii) segmentation of NM clusters within the somata. The cell bodies of the DNs were segmented based on the nickel map for the sections stained for TH with nickel enhancement. For each animal we manually labeled three to five neurons and a region containing no neurons. The classifier was trained on these labels in five ROIs using the feature maps including information on intensities (raw image, raw image smoothed with gaussian kernels (*σ* = 1, 3.5 px)), edges (Laplacian of Gaussian, applied on smoothed raw image with gaussian kernel, *σ* = 10 px, Gaussian gradient magnitude (3.5, 10 px), difference of Gaussians (3.5, 10 px)), and textures (structure tensor eigenvalues (5 px) and Hessian of a Gaussian eigenvalues (5 px)).

The NM clusters were identified as sulfur-rich regions on the sulfur maps in both unstained and stained sections. NM is known to contain an elevated concentration of sulfur due to the presence of sulfur-rich compounds, such as benzothiazine rings and cysteine residues [16]). For all animals except for the youngest, in which no NM clusters were detected, three to five NM clusters and regions with no NM clusters were labeled manually. The classifier was trained on these labels in eight ROIs. For the stained sections, the sulfur masks were additionally constrained by the nickel mask to reduce artifacts arising from sulfur-rich structures other than intracellular NM. For the youngest animal (0.1 y), which did not show any NM, the contrast of the sulfur maps was not high enough to derive segmentation. For this animal, only the nickel based segmentation on TH-stained section was used. This should not introduce any bias to the lifespan trajectory of iron concentrations in NM clusters, because the iron was very homogeneously distributed across the tissue and within neurons (SD(Fe) *<* 20 μg g^−1^), thus the choice of mask did not impact the obtained cellular concentrations.

Iron measurements were corrected for the contribution of iron present in the embedding medium by subtracting the mean iron concentration measured in regions containing only the medium (i.e. no tissue), from the cellular iron concentrations per ROI.

In order to facilitate data comparison for valuable data like element quantification using μXRF, we want to emphasize that it is necessary to provide reference data or an example of delineation of the measured areas, see Fig. 1, Suppl. Fig.8.

We estimated a potential systematic bias of the iron concentrations due to (i) the immunohistochemical staining, (ii) the intra-individual anatomical variability in iron concentrations and (iii) the variability of iron concentrations due to the neuronal segmentation by performing Welch two-sample-t-tests, see Suppl. Fig. 10. A p-value of ≤ 0.001 was considered significant. Additionally, we computed the relative difference of medians 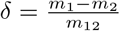, where *m*_1_ and *m*_2_ are the medians of the two sets and *m*_12_ is the median of the joint set.

### 4.6 Bayesian modeling of lifespan iron accumulation

To characterize the lifespan accumulation of iron within cells and in SN tissue, we modeled the age dependence of iron markers derived from qMRI and μXRF using the exponential saturation function proposed by Hallgren and Sourander for brain iron accumulation [35]:

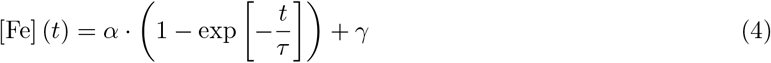

where *t* is the age in years, *α* is the amplitude of overall changes in iron concentration throughout the lifespan, *τ* is the time constant of the exponential saturation, and the *γ* represents the iron concentrations at birth.

To fully exploit the sparsely sampled data across the lifespan, particularly in the μXRF dataset, we used a Bayesian approach. We used a Gaussian likelihood function with mean [Fe] to model the iron concentration as a function of age *t*. Assuming weakly informed prior distributions of model parameters *α, τ* and *γ*, we estimated their posterior distributions given experimental data. The dependent variable was the iron marker (median of mean cellular iron concentrations in DN or median 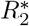 in SN) as a function of the age *t*. The independent variables were parameters *α, τ* and *γ* as well as *σ*, the standard deviation of iron concentrations (or 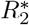 values). The standard deviation of iron concentrations was modeled as a normal distribution. For the modeling the standard deviation of the 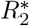 trajectory, we modeled a linearly increasing *σ* = *u* + *v* · *t*, where *u* and *v* were halfnormal distributions with standard deviation of 10 y, to model the increasing variability of 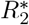 with age *t*, see Suppl. Fig.12a. Broad priors were chosen for the modeling of the iron concentration: parameter *α* (uniformly distributed between 100 and 1000 μg g^−1^, order of magnitude from previous data [36]) and the intercept parameter *γ* (uniform between 0 and 100 μg g^−1^, order of magnitude as reported in [35]). Similar broad priors were chosen for the modeling of 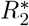 (*α*: uniform from 10 to 1000 s^−1^ and *γ*: 10 and 100 s^−1^). For the time constant *τ*, we used the same Gaussian distribution as the prior for both models, with a mean of 16 years and a standard deviation of 25 years informed by time constants reported across subcortical and cortical regions in humans [35]).

The Bayesian model was implemented in PyMC v.5.16.1 in python and visualized using Arviz v.0.18.0. Four thousand samples were drawn, while the first 2000 were neglected for tuning of each of the 4 chains.

Trace plots were used as quality checks for Markov Chain Monte Carlo (MCMC) chain convergence, see Suppl. Fig. 13 and 14. Bayes factors for the model of cellular iron concentrations in chimpanzee DN were: BF_*α*=409_ = 8.1, 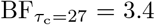, BF_c=45_ = 4.6. Bayes factors for the model of average iron concentrations in chimpanzee SN were: BF_*α*=75_ = 20, 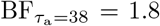, BF_c=15_ = 16.6. A summary of posterior distribution of variables with a 68% mass of credible interval is shown in Suppl. Fig. 12. To compare the lifespan trajectories of cellular iron in DNs and average tissue iron in the SN, as well as cellular iron accumulation in DN between humans and chimpanzees, we compared the posterior distributions of the time constant parameter *τ*, obtained for cells and tissue and for both species. We evaluated the contrast of posterior distributions, i.e. the distribution of the difference of the different posterior distributions [74]. The code for the Bayesian modeling will be provided upon acceptance.

### 4.7 Neuromelanin quantification from optical microscopy

NM concentrations of DN were quantified in 3 chimpanzees and 2 humans (sample 2 and 3 described in [41]) using optical microscopy and the published extinction coefficients of NM [57]. The absorbance of NM was shown to depend linearly on the NM concentration, independent of its iron load [75, 76]. NM is composed of the two chromophores pheomelanin (mass fraction within NM 23 %) and eumelanin (77 %) [56].

First, we calculated optical density OD maps from brightfield microscopy by taking the natural logarithm of the microscopy intensity of the blue channel *I* normalized by the intensity in a region without NM *I*_0_: OD = −log(*I/I*_0_). Second, we computed the chromophore concentration using the reported wavelength-dependent extinction coefficients *c*_*λ*_ from the literature at the spectral density maximum of the camera (AxioScan rgb color camera 705, Zeiss, Jena) and section thickness *d* = 10 μm (Lambert-Beer law, [57]):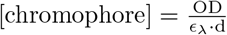. Third, we calculated cellular NM concentrations by dividing the resulting chromophore concentrations by the chromophore-NM ratio. This scaling factor was determined from the chromophore-sulfur ratio using the semi-quantitative sulfur maps from μXRF together with the published sulfur-NM ratio of 0.0256 [77, 78]. A chromophore-sulfur ratio of 15.5 was determined as the mean of slopes of the linear relation of chromophore to sulfur concentrations in the 44-year-old chimpanzee and 86-year-old human (Suppl. Fig. 11a). We assume that the composition of NM including lipids, proteins and chromophores, is similar to that of NM extracted in the protocol of [22, 79]. We segmented individual NM clusters in DN using a trained classifier on the optical microscopy.

The iron and NM concentrations both vary significantly across age and species (separate ANOVA for both iron and NM, p *<* 0.001). Neither iron nor NM concentrations showed significant variation across adjacent sections for one individual, nor did their slope differ (61-year old human, ANOVA iron: p = 0.249, NM: p = 0.646, ANCOVA: iron: NM *×* section p = 0.006). There was no evidence for a difference in the slope of iron and NM concentrations in the adolescent and adult chimpanzees (ANCOVA: p = 0.720). There was also no evidence that the slope of iron and NM concentration in the younger and older human sample differed (ANCOVA: NM *×* age interaction p = 0.851). However, the sparse data showed evidence that the slope of iron and NM differed significantly between the species. This resulted from a generalized linear model and was supported by a Bayesian ANCOVA that identified NM and NM *×* species terms as the most probable predictors of the cellular iron concentration (dependent variable iron concentration, covariate NM, fixed factors species and age, including individual as a random factor: NM and species vs. NM-only: BF_10_ = 0.568, other models tested: null model, species only, age only, age *×* species, age *×* species *×* NM). ANOVA, ANCOVA, correlation analysis, generalized linear model and Bayesian ANCOVA were performed in JASP 0.95.1.

### 4.8 Iron-binding model

We assumed a chemical equilibrium between labile iron in the cytosol Fe^labile^ and iron bound to two binding sites in NM – one high-affinity binding and another low-affinity binding site:

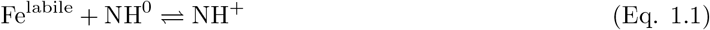

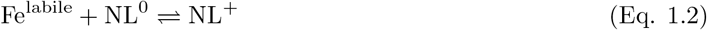

Where NH^0^ and NH^+^ are the vacant and occupied high-affinity binding sites in NM, respectively, and NL^0^ and NL^+^ are the vacant and occupied low-affinity binding sites in NM, respectively.

In chemical equilibrium between labile iron and the high- and low-affinity binding sites, the following equation holds:

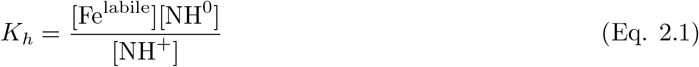

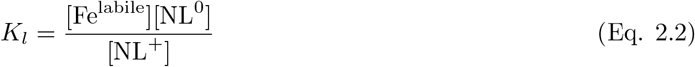

We introduce the dissociation constants for high- and low-affinity binding sites, *K*_*h*_ and *K*_*l*_, the concentrations of labile iron [Fe^labile^], vacant high-affinity binding sites [NH^0^], occupied high-affinity binding sites [NH^+^], vacant low-affinity binding sites [NL^0^] and occupied low-affinity binding sites [NL^+^] at equilibrium.

The total concentration of iron is given by:

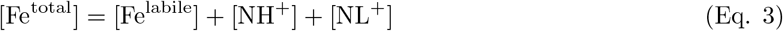

We assume that the total cellular concentration of NM, [NM], is known from optical absorption measurements, and the total iron concentration, [Fe^total^], is known from XRF measurements. Using this information, we aimed to estimate the concentration of labile, toxic iron in the cytosol, [Fe^labile^].

We denote the fraction of occupied high-affinity binding sites as *h* and the fraction of occupied low-affinity binding sites as *l*. The maximum possible molar concentrations of iron in the high- and low-affinity binding sites are H and L, respectively. These four parameter were previously determined for extracts of human NM ([22]) and are summarized in the Table 4.

**Table 4.**
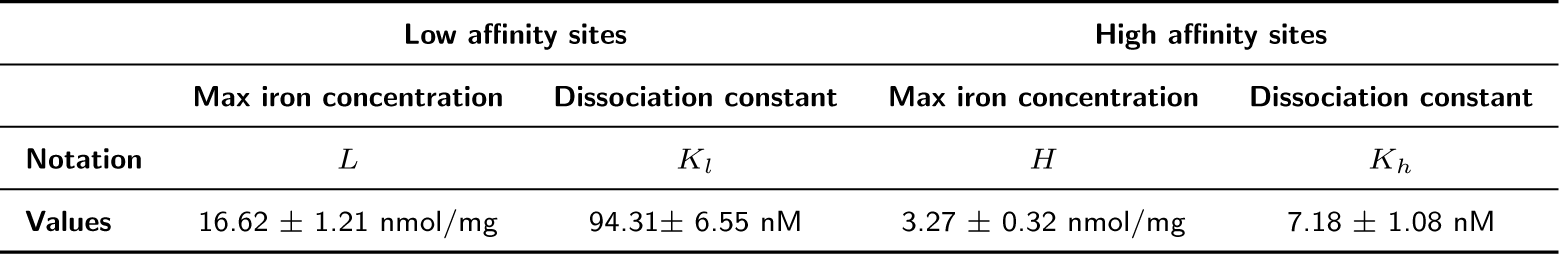
Iron-binding characteristics of two iron binding sites in NM, reproduced from [22] (Table 1).

Using these definitions, Eqs. 2 and 3 can be rewritten in terms of *N, H, L, h*, and *l* as follows:

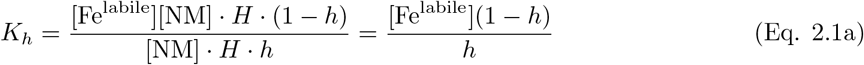

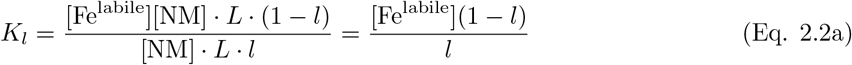

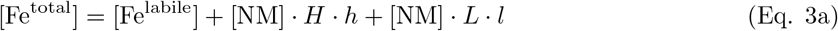

Let us introduce new variables 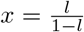 and 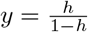, so that 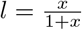 and 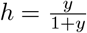 . Then, the system of Eqs. 2a, 2b and 3a can be rewritten as a system of three equations with three unknowns:

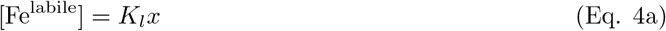

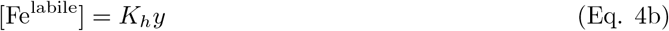

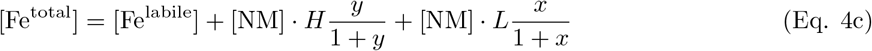

By combining Eqs. 4a and 4b, we obtain the relationship between *x* and *y*:

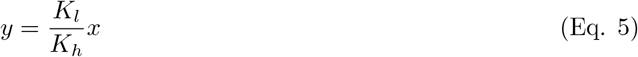

Dividing Eq. 4c by the maximum possible iron concentration bound to NM, [Fe^max^] = [NM] · *H* +[NM] · *L*, we derive a dimensionless equation:

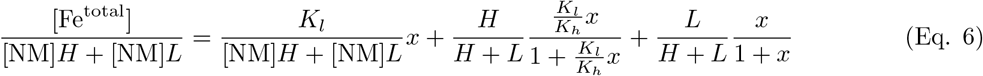

Denoting 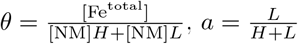, and 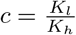, Eq. 6 can be simplified as follows:

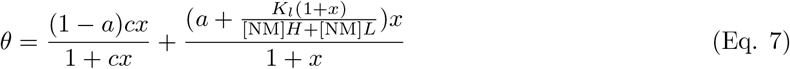

For the maximum possible NM concentration of 10^6^ mg/L, and a saturation ratio of low affinity binding sites *l* not too close to 1, the term

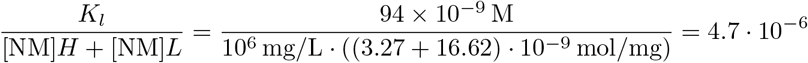

is negligible compared to

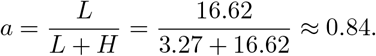

Thus, Eq. 7 simplifies to a quadratic equation:

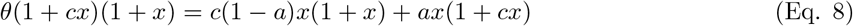

Expanding Eq. 8, we obtain:

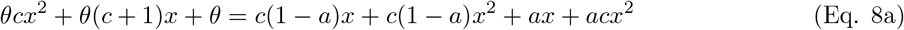

Simplifying further:

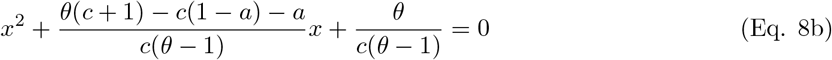

Denoting 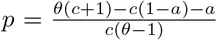 and 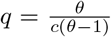, we solve Eq. 8b, choosing only the physically meaningful positive solution:

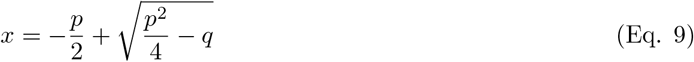

The cellular concentration of labile iron can then be determined by substituting Eq.9 into Eq. 4a:

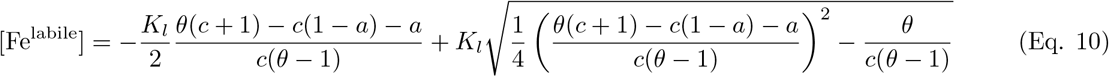

Note that [Fe^labile^] depends only on the dimensionless parameter 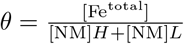, representing the NM-iron-saturation fraction, and the constants *H, L, K*_*l*_, and *K*_*h*_, which have been determined in independent experiments (Table 1, [22]). Estimated labile iron, low and high affinity bound iron as well as total iron as a function of *θ* is visualized in Suppl. Fig. 15.

## 5 Data availability

The elemental maps from μXRF including the segmentation masks used in this study will be made available upon publication. Other relevant data is available from the authors on reasonable request.

## 6 Acknowledgments

This is an EU Joint Program - Neurodegenerative Disease Research (JPND) project. The project is supported through the following funding organizations under the aegis of JPND - https://neurodegenerationresearch.eu, the Federal Ministry of Education and Research (BMBF) under support code 01ED2210, and the Federal Ministry of Research, Technology and Space (BMFTR) under grant number 01ED2508. The project has received funding from the Deutsche Forschungsgemeinschaft (DFG, German Research Foundation) – project no. 347592254 (WE 5046/4-2 and KI 1337/2-2). and was funded by the Max Planck Society under the inter-institutional funds of the president of the Max Planck Society for the Hominoid Brain Connectomics (EBC) Project (M.IF.NEPF8103 and M.IF.EVAN8103). Felix Büttner was supported by the International Max Planck Research School on Cognitive Neuroimaging. We acknowledge DESY (Hamburg, Germany), a member of the Helmholtz Association HGF, for the provision of experimental facilities. Parts of this research were carried out at PETRA III and we would like to thank K. V. Falch and J. Garrevoet for assistance in using the microprobe at beamline P06. Beamtime was allocated for proposal I-20211534. This research was supported in part through the Maxwell computational resources operated at DESY. We thank Kerrin Pine for the implementation of MRI sequences used in the study and Kay Double, Sylvain Bohic and Harald Möller for the fruitful discussion. Furthermore, we would like to thank Alex Crabbe and Siawoosh Mohammadi for their support with the experiments at beamline P06 and Anne Schmieder for her help with segmenting the SN in MRI volumes.

## 7 EBC Consortium

https://www.eva.mpg.de/ecology/projects-and-research-groups/evolution-of-brain-connectivity/

Bala Amarasekaran (Tacugama Chimpanzee Sanctuary, Freetown, Sierra Leone)

Caroline Asiimwe (Budongo Conservation Field Station, Uganda)

Daniel Aschoff (German Primate Center, Göttingen, Germany)

Felix Büttner (Max Planck Institute for Human Cognitive and Brain Sciences, Leipzig, Germany)

Malte Brammerloh (Microstructure Mapping Lab, Lausanne, Switzerland)

Dennis Brückner (DESY, Hamburg, Germany)

Penelope Carlier (Tai Chimpanzee Project, CSRS, Abidjan, Côte d’Ivoire)

Julian Chantrey (Veterinary Pathology and Preclinical Sciences, University of Liverpool, UK)

Catherine Crockford (Max Planck Institute for Evolutionary Anthropology, Leipzig, Germany)

Tobias Deschner (Institute for Cognitive Sciences, University of Osnabrück, Germany; Ozouga, Loango Chimpanzee Project, Loango National Park, Gabon, Max Planck Institute for Evolutionary Anthropology, Leipzig, Germany)

Ariane, Düx (Robert Koch Institute, Berlin, Germany; Helmholtz Institute for One Health, Helmholtz Centre for Infection Research, Greifswald, Germany)

Geraldine Escoubas (Max Planck Institute for Evolutionary Anthropology, Leipzig, Germany; Loango Chimpanzee Project, Loango National Park, Gabon)

Gerald Falkenberg (DESY, Hamburg, Germany)

Pawel Fedurek (School of Psychology, University of Stirling, UK; Budongo Conservation Field Station, Masindi, Uganda)

Alejandra Romero Forero (Tacugama Chimpanzee Sanctuary, Freetown, Sierra Leone)

Angela D. Friederici (Max Planck Institute for Human Cognitive and Brain Sciences, Leipzig, Germany)

Zoro B. GoneBi (Department of Bioscience, University Felix Houphouet-Boigny, Abidjan, Côte d’Ivoire; Tai Chimpanzee Project, CSRS, Abidjan, Côte d’Ivoire)

Tobias Gräßle (Helmholtz Institute for One Health, Helmholtz Centre for Infection Research, Greifswald, Germany; Robert Koch Institute, Berlin, Germany)

Philipp Gunz (Max Planck Institute for Evolutionary Anthropology, Leipzig, Germany) Daniel Hanus (Zoo Leipzig, Leipzig, Germany)

Jenny E. Jaffe (Robert Koch Institute, Berlin, Germany; Tai Chimpanzee Project, CSRS, Abidjan, Côte d’Ivoire)

Carsten Jäger (Max Planck Institute for Human Cognitive and Brain Sciences, Leipzig, Germany)

Anna Jauch (Paul Flechsig Institute, Center of Neuropathology and Brain Research, Leipzig University, Leipzig, Germany)

Evgeniya Kirilina (Max Planck Institute for Human Cognitive and Brain Sciences, Leipzig, Germany)

Kathrin Kopp (Max Planck Institute for Evolutionary Anthropology, Leipzig, Germany)

Fabian H. Leendertz (Helmholtz Institute for One Health, Helmholtz Centre for Infection Research, Greifswald, Germany; Robert Koch Institute, Berlin, Germany)

Ilona Lipp (Max Planck Institute for Human Cognitive and Brain Sciences, Leipzig, Germany)

Matyas Liptovszky (Twycross Zoo, UK)

Patrice Makouloutou Nzassi (Institut de Recherche en Ecologie Tropicale, Libreville, Gabon)

Kerstin Mätz-Rensing (German Primate Center, Göttingen, Germany)

Matthew McLennan (Bulindi Chimpanzee and Community Project, School of Social Sciences, Oxford Brooks University, UK)

Richard McElreath (Max Planck Institute for Evolutionary Anthropology, Leipzig, Germany)

Sophie Moittie (Twycross Zoo, UK) Torsten Møller (Kolmarden Zoo, Sweden)

Markus Morawski (Paul Flechsig Institute, Center of Neuropathology and Brain Research, Leipzig University, Leipzig, Germany)

Karin Olofsson-Sannö (National Veterinary Institute, Uppsala, Sweden)

Simone Pika (University of Osnabrück, Osnabrück, Germany; Ozouga Chimpanzee Project, Loango National Park, Gabon)

Kerrin Pine (Max Planck Institute for Human Cognitive and Brain Sciences, Leipzig, Germany)

Andrea Pizarro (Tacugama Chimpanzee Sanctuary, Freetown, Sierra Leone)

Kamilla Pleh (Robert Koch Institut, Berlin, Germany: Tai Chimpanzee Project, CSRS, Abidjan, Côte d’Ivoire)

Tilo Reinert (Paul Flechsig Institute, Center of Neuropathology and Brain Research, Leipzig University, Leipzig, Germany)

Jessica Rendel (Twycross Zoo, UK)

Liran Samuni (Human Evolutionary Biology, Harvard University, Cambridge, USA; Tai Chimpanzee Project, CSRS, Abidjan, Côte d’Ivoire)

Mark Stidworthy (International Zoo Veterinary Group, Keighley, UK)

Lara Southern (Institute for Cognitive Sciences, University of Osnabrück, Germany; Ozouga, Loango Chimpanzee Project, Loango National Park, Gabon)

Tanguy Tanga (Institut de Recherche en Ecologie Tropicale, Libreville, Gabon; Ozouga, Loango Chimpanzee Project, Loango National Park, Gabon)

Reiner Ulrich (Institute for Veterinary Pathology, Leipzig University) Steve Unwin (Wildlife Health Australia, Sydney, Australia) Sue Walker (Chester Zoo, UK)

Nikolaus Weiskopf (Max Planck Institute for Human Cognitive and Brain Sciences, Leipzig, Germany) Roman M. Wittig (Max Planck Institute for Evolutionary Anthropology, Leipzig, Germany)

Kim Wood (Welsh Mountain Zoo, UK)

Klaus Zuberbuehler (Institute of Biology, University of Neuchatel, Switzerland; Budongo Conservation Field Station, Masindi, Uganda)

## 8 Author contributions

Conceptualization: F.B., E.K., N.W., M.M., Methodology: F.B., E.K., N.W., T.R., G.F., D.B., M.B., C.J., M.M., C.C., R.W., R.M., T.D., Formal analysis: F.B., E.K., M.B., D.B., Investigation: F.B., E.K., T.R., I.L., M.B., G.F., D.B., C.J., M.M., Writing-original draft: F.B., E.K., Writing-review and editing: all authors, Visualization: F.B., E.K., Supervision: E.K. and N.W., Funding acquisition: N.W., E.K., R.W., C.C., All authors contributed to the paper and approved the submitted version.

