## Supplementary Information for "Single-cell quantification of the iron-neuromelanin balance in dopaminergic neurons across the lifespan"

### 9 Supplementary Information

#### 9.1 SN tissue iron quantification across the lifespan using qMRI

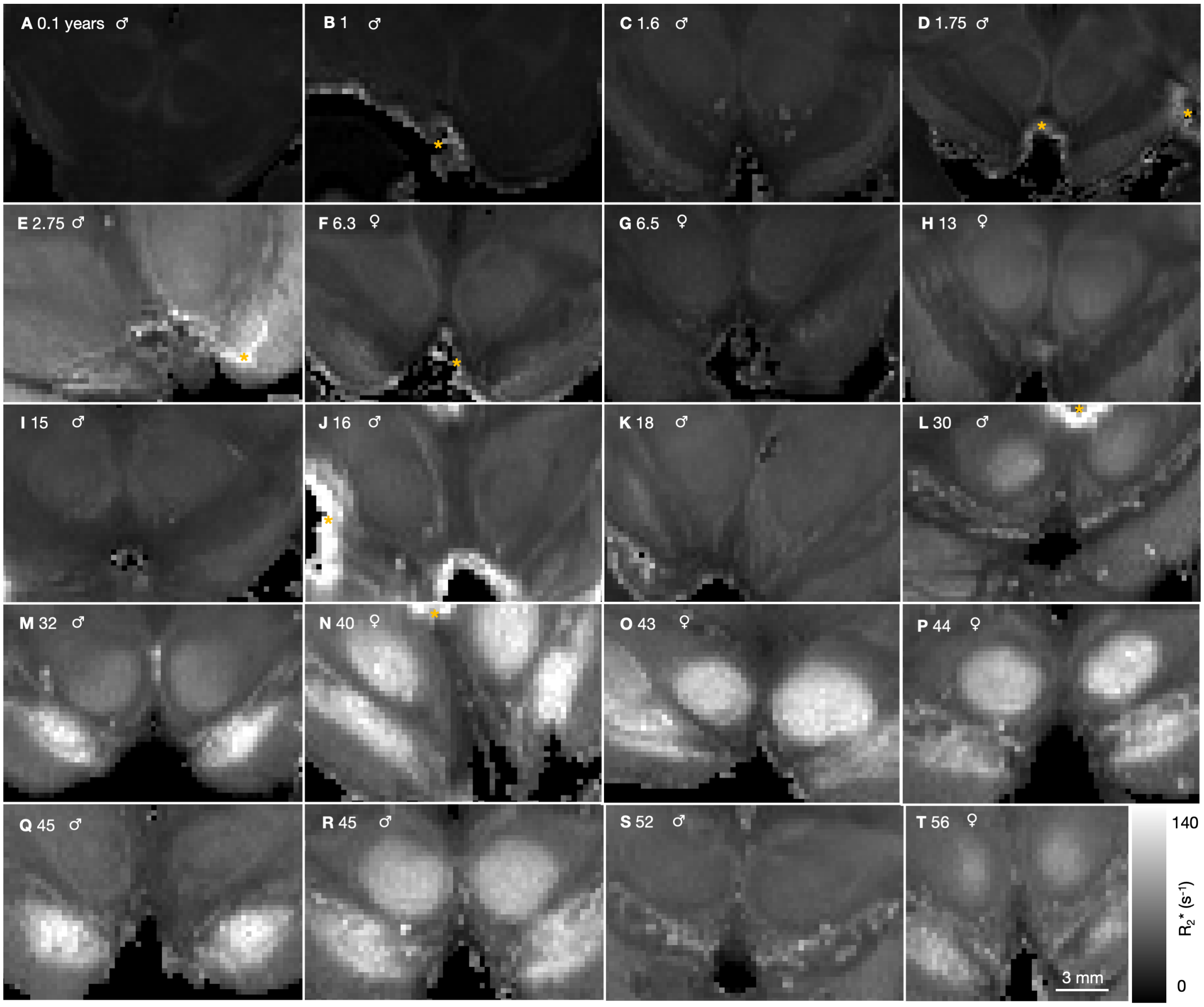


**Figure 6:** Quantitative maps of the effective transverse relaxation rate $R_2^*$ across the chimpanzee lifespan. Zoom-in into the midbrain (coronal slice) including the nucleus ruber and the SN are shown. Animals are arranged in the order of increasing age, from 0.1 to 56 years. The maps are equally color-scaled for all ages, enabling easy comparison. Both, the overall $R_2^*$, and the $R_2^*$ contrast within the SN increased with age showing hyperintense substructures within the SN from an age of 13 years on, corresponding to SNpc with high density of NM-pigmented neurons. Yellow stars indicate regions with artifacts resulting from tissue damage or air bubbles within the samples that were therefore excluded from the analysis.

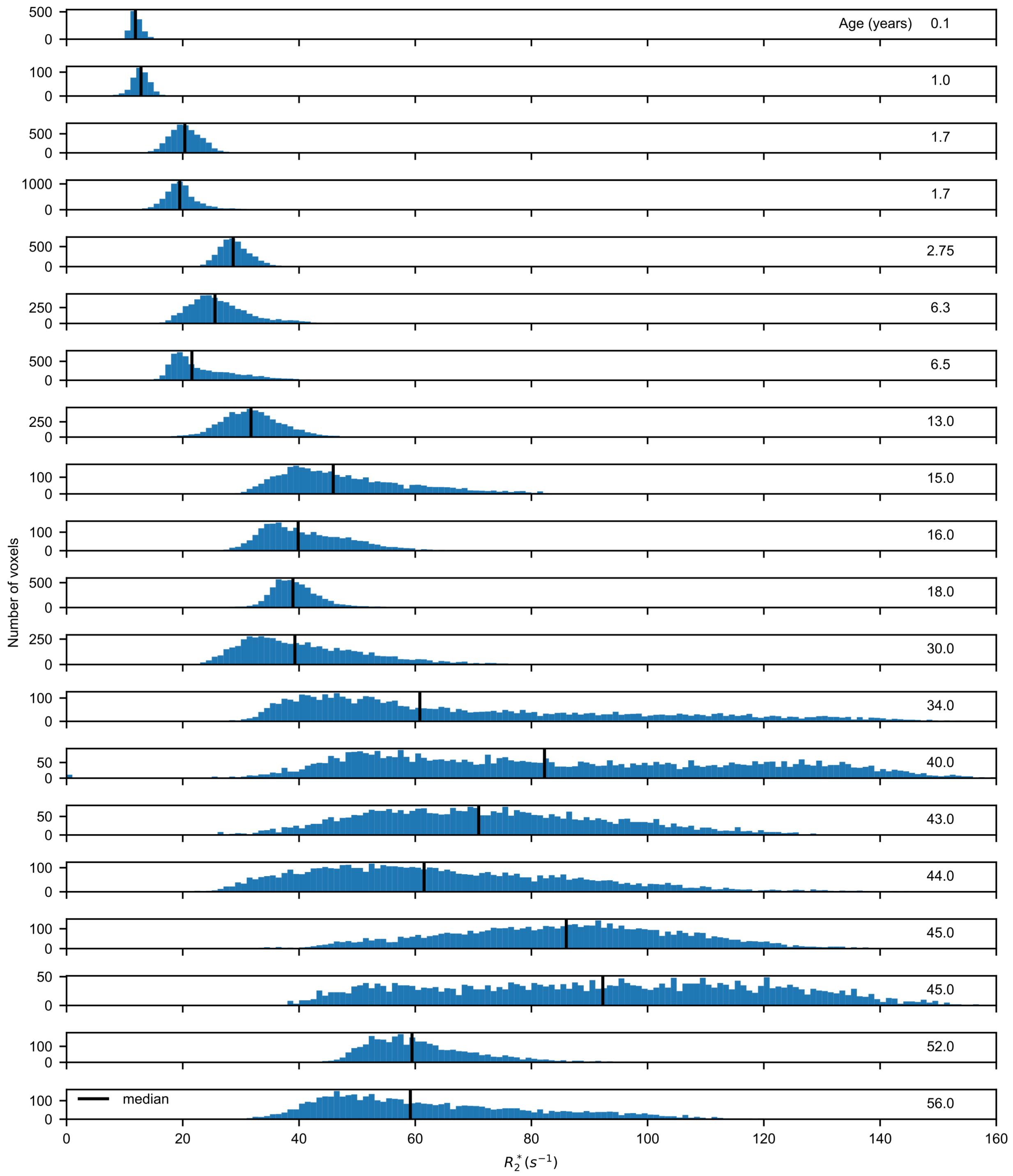


**Figure 7:** Histograms of $R_2^*$ in the SN of 20 chimpanzees. The median $R_2^*$ increases across the lifespan (solid black line) and was used for modeling the lifespan trajectory in Fig. 3. For the elderly adults (> 30 years) the distribution showed a larger variance.

#### 9.2 Tissue samples

| Case | Age(y) | Sex | Subspecies | Site | PMI (h) | Cause of death |
|---|---|---|---|---|---|---|
| A | 0.1 | M | schweinfurthii | wild | 4 | con-specific aggression |
| B | 1±1 | M | troglodytes | wild | 4.5 | infanticide |
| C | 1.6 | M | verus | sanctuary | <16 | con-specific aggression |
| D | 1.75 | M | verus | wild | 24 | Bcbva infection |
| E | 2.75 | M | verus | wild | 3.5 | starvation |
| F | 6.3 | F | verus | wild | 4 | con-specific aggression |
| G | 6.5±1 | F | verus | sanctuary | 5 | unknown |
| H | 13±3 | F | schweinfurthii | wild | 15 | inter-specific aggression |
| I | 15 | M | verus | wild | <12 | Bcbva infection |
| J | 16 | M | verus | wild | <16 | Bcbva infection |
| K | 18±3 | M | troglodytes | wild | 11 | con-specific aggression |
| L | 30±5 | M | schweinfurthii | wild | 12 | inter-specific aggression |
| M | 32 | M | verus | zoo | 6 | urolithiasis |
| N | 40±5 | F | verus | wild | <24 | chronic renal disease |
| * | 41 | F | verus | zoo | 4 | weakness, pneumonia |
| O | 43 | F | schweinfurthii | zoo | 1 | unknown |
| P | 44 | F | mixed | zoo | 1 | cardiovascular diseases |
| Q | 45±5 | M | verus | wild | 14.5 | leopard attack |
| R | 45 | M | unknown | zoo | <24 | cardiovascular diseases |
| * | 47 | F | verus | zoo | 4 | abdominal tumours |
| S | 52 | M | verus | zoo | <24 | epileptic episode |
| T | 56±5 | F | verus | wild | 2.5 | leopard attack |

**Table 5:** Tissue information for the *post mortem* brain samples from 20 chimpanzees (*Pan troglodytes*) used for qMRI. The *post mortem* interval (PMI) denotes the time span between death and brain fixation. For the subset of five chimpanzees (gray rows), the midbrain was processed for histology and XRF corresponding to case numbers 1,2,3,4 and 5. Bcbva: Bacillus cereus biovar anthracis. * Two individuals were excluded from the analysis due to imaging artifacts in the SN.

895 ## 9.3 Cell-specific iron quantification using X-ray fluorescence

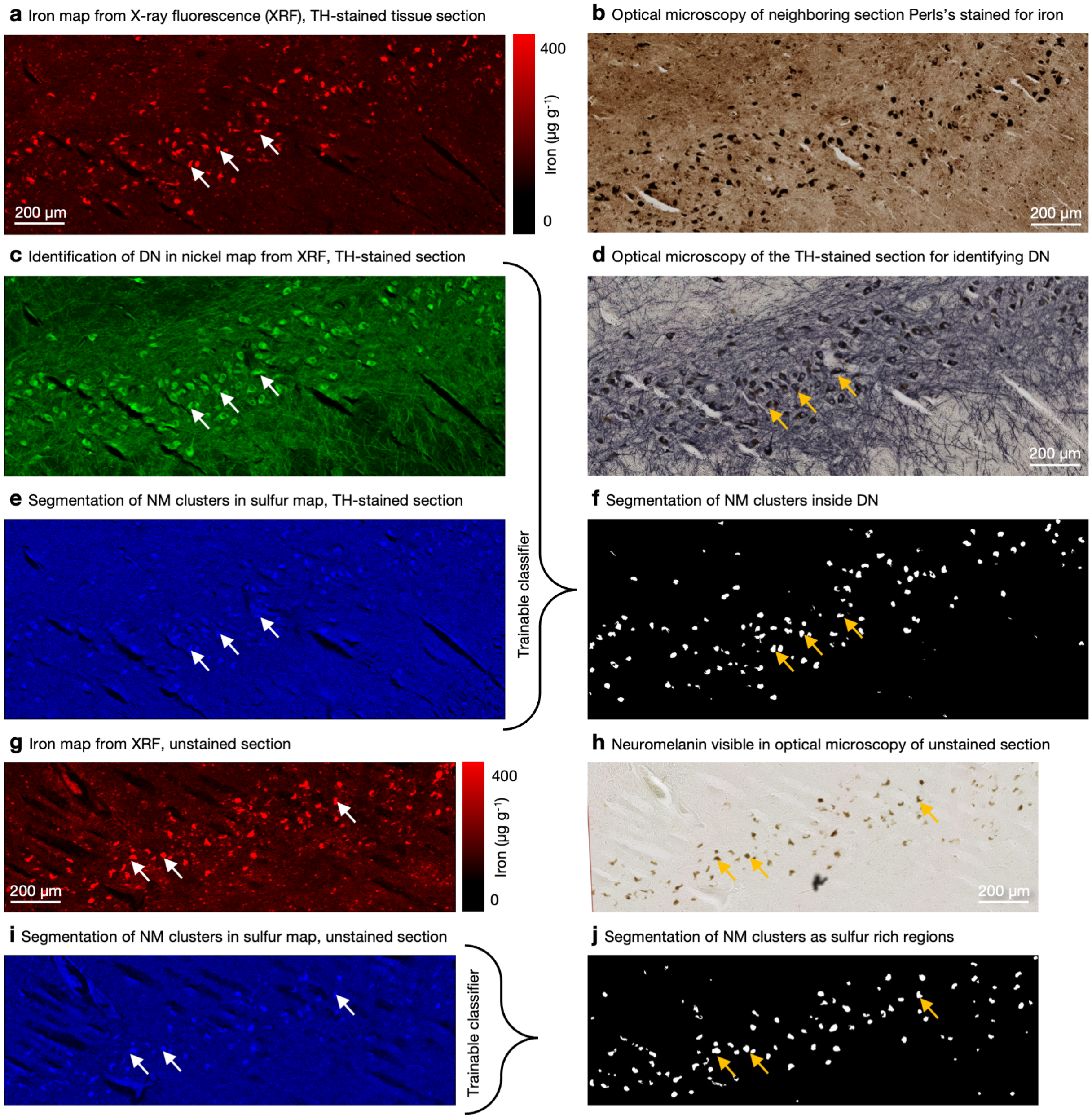


**Figure 8:** Quantitative element mapping by µXRF (left column) that was used for cellular iron quantification is compared to conventional immunohistochemical staining and cellular segmentation of NM clusters (right column). **a & b** The quantitative iron map mirrored the increased iron load of DN – as seen in a classical Perls's stain for iron of the neighboring section – but provided quantitative information about cellular iron concentrations. **c, d, e & f** Tissue section stained for tyrosine hydroxylase (TH) were used to identify NM clusters (using the sulfur map from XRF) restricted within the DN cell bodies (identified using the nickel map, as the TH-staining was enhanced by nickel). **g, h, i & j** In unstained tissue sections, sulfur maps were used to segment NM clusters and measure the iron concentration wihtin NM and control for potential elemental redistribution during the staining process. To trace the lifespan trajectory of cellular iron concentrations of DN, data from both unstained and stained tissue was pooled in order to increase the number of analyzed cells and therefore statistical power. All subfigures show data from sample 5 (44 years, see Table 3). The arrows point to the same NM clusters, showing that regions of high iron concentration are overlapping with regions of high sulfur concentration and darker regions in optical microscopy as seen in the unstained section (h). For the stained section, we observe that these iron-rich regions are located inside DNs, as labeled with nickel (c).

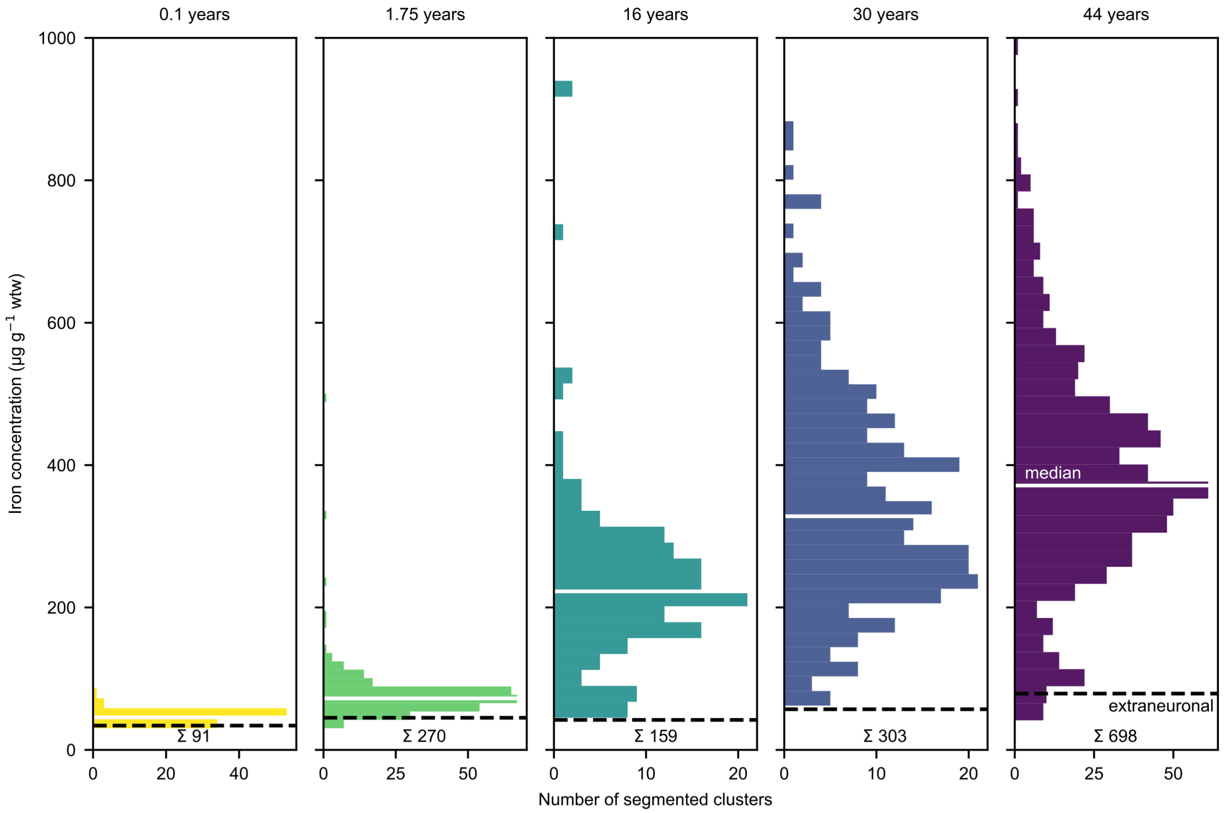


**Figure 9:** The iron concentration in the NM clusters of DN measured by µXRF combined with Ni-enhanced immunohistochemistry showed a large increase across the lifespan. The histograms show the large number of automatically segmented clusters for the pooled unstained and TH-stained sections including two ROIs for each of the two sections per brain. Two additional TH-stained sections are included for the oldest animal. The solid white lines indicate the median NM-iron concentration, the dashed lines show the iron concentration outside the NM mask. For the two young animals, 0.1 and 1.75 years, the NM-iron concentration is in the range of the background iron concentration, while NM-iron is much higher and wide spread for the older animals. The total number of analyzed NM clusters is provided at the bottom ($\Sigma$).

#### 9.4 Sources of error and variability in cellular iron quantification

The age-related increase in cellular iron concentration reported in this manuscript is prone to several systematic and random measurement errors. As shown below, the biological effect of interest, which is the increase of iron concentration across the lifespan, is larger than the errors resulting from the observed anatomical variability, the effect of the histological treatment, the measurement error of µXRF and the influence of the segmentation of NM in DN, Fig. 10. The increase with age was highly significant ($p \leq 0.001$) between succeeding individuals. The iron concentration increased by 41 % from 0.1 to 1.75 years, by 182 % from 1.75 to 16 years, by 38 % from 16 to 30 years and by 12 % from 30 to 44 years.

*Measurement error of the µXRF*: Random and systematic measurement errors of the iron quantification experiment contributed to the variance in iron concentration. The systematic error of the µXRF calibration was at least 10%, based on the absolute precision of used calibration standard. Note that this error impact the data of all animal and humans, so it does not impact the measured time constants of life span dynamics. The random error of cellular XRF measurements due to the random nature of x-ray-photon-absorption and -emission processes can be estimated using Poisson statistics, given that on average around 50 photons per cell were detected. This result in random Poissonian noise with a standard deviation of 14%.

*Systematic bias due to immunohistochemical staining*: We tested if the staining process influenced the iron concentration in the NM clusters of DN. While the iron concentration in stained tissue was slightly higher for sample 2 (relative difference +21%), the iron concentration in stained tissue was lower in sample 3 ($p < 0.001$, −38%). However, as these sections were approx. 100 µm apart, we can not exclude that this difference also includes an effect of iron variability due to the anatomical location, Fig. 10a.

*Variability due to neuronal segmentation*: The variability of the median iron concentration of cell populations due to the segmentation using a trainable classifier was assessed by training the classifier by a second rater. It was found to be less than 14%, Fig. 10c, d.

*Intra-subject anatomical variability*: We measured the iron concentration in NM clusters of two different ROIs for neighboring sections. The concentrations in the two different ROIs differed not significantly ($p = 0.54$). The anatomical variability was about 3 % (Fig. 10b) and originated probably from the spatial iron concentration gradient within the SN.

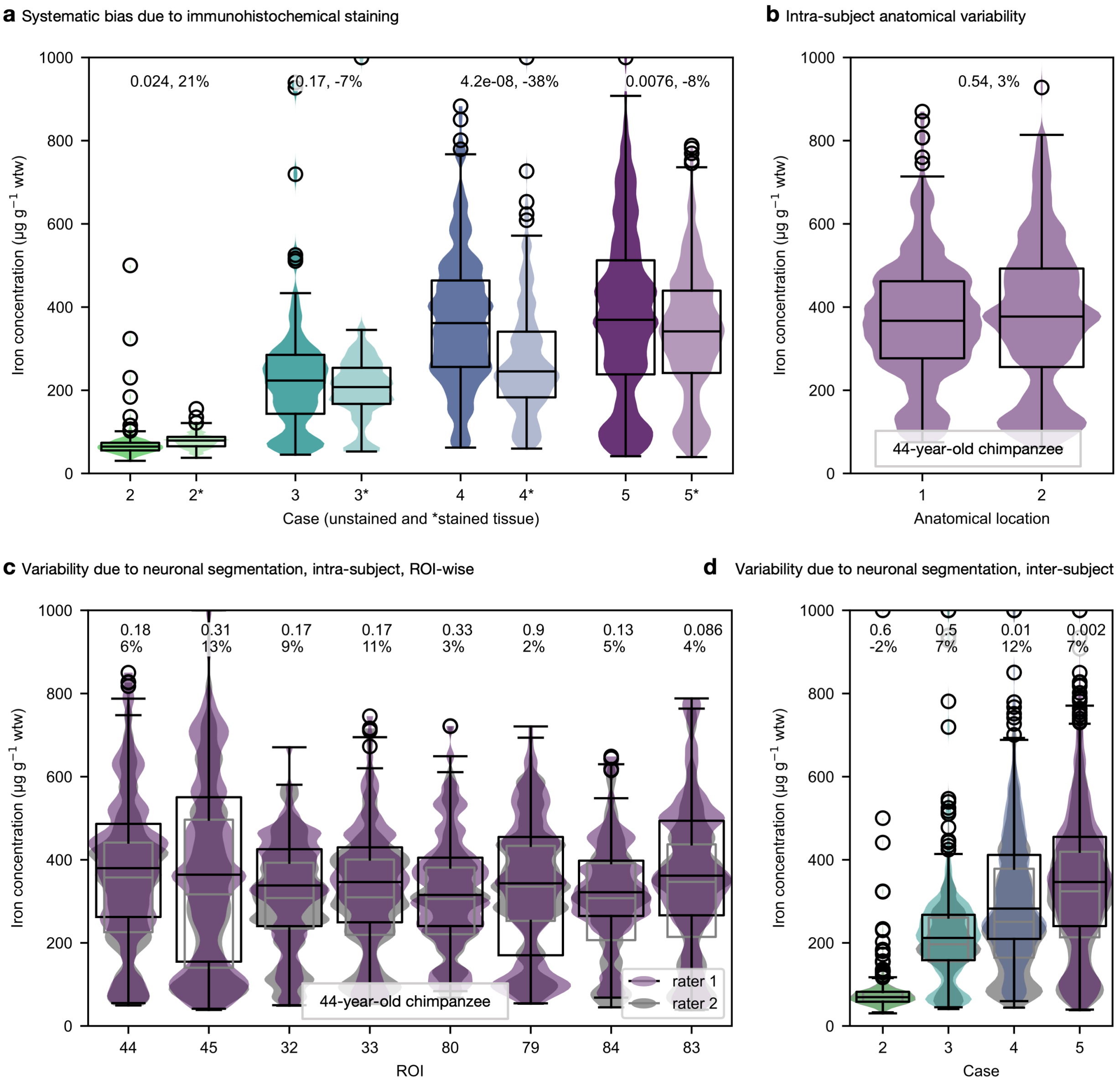


**Figure 10: Sources of variability in the cellular iron quantification using XRF. a** The iron concentration of individual NM clusters for unstained (intense color) and stained (* faint color) tissue showed similar distributions. **b** The iron concentration of individual NM clusters for two different ROIs pooled over three adjacent sections of case 5 did not differ significantly. The medians showed a relative difference of about $3\,\%$, indicating little anatomical variability. **c** Labeling of the NM clusters and neurons to train the classifier was performed by a second rater (gray violin- and boxplots) to assess the influence of the segmentation on the variability of iron concentrations. The variation of median iron concentration in 8 different sections (stained and unstained) was below $14\,\%$. ROIs 45 and 80 were used for labeling. **d** Similar low variability was found across all cases. Violin plots show the density function, the box plots indicate the interquartile range and the medians. For visualisation the data is clipped at $1000\,\mu g\,g^{-1}$. P-values and relative differences are displayed above the corresponding data.

#### 9.5 Neuromelanin quantification in dopaminergic neurons using optical microscopy

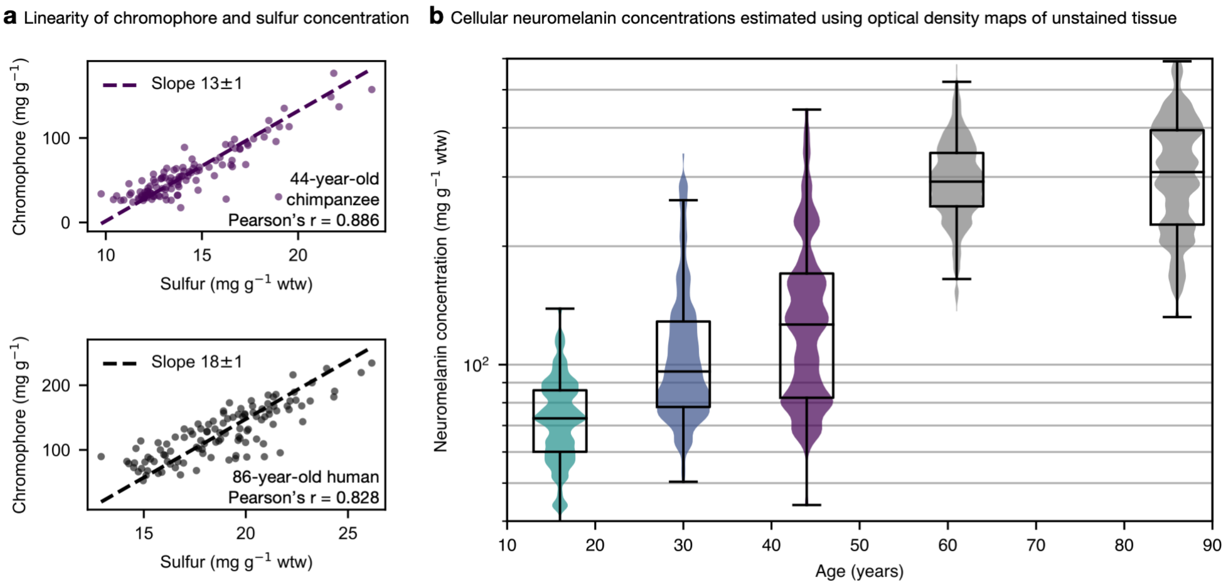


**Figure 11:** Neuromelanin concentrations in DN are increasing with age. **a** The linearity of the dependence of sulfur concentrations obtained using XRF and chromophore concentrations determined from optical microscopy in individual neurons of the 44-year-old chimpanzee and the 86-year-old human supports the assumption that the optical density of DN is mainly modulated by their NM concentration. The slope of the orthogonal distance regression is in mg chromophore per mg sulfur. **b** The NM concentration in DN is increasing with age in chimpanzees (colored violins) and humans (gray violins).

#### 9.6 Bayesian modeling of lifespan iron trajectories

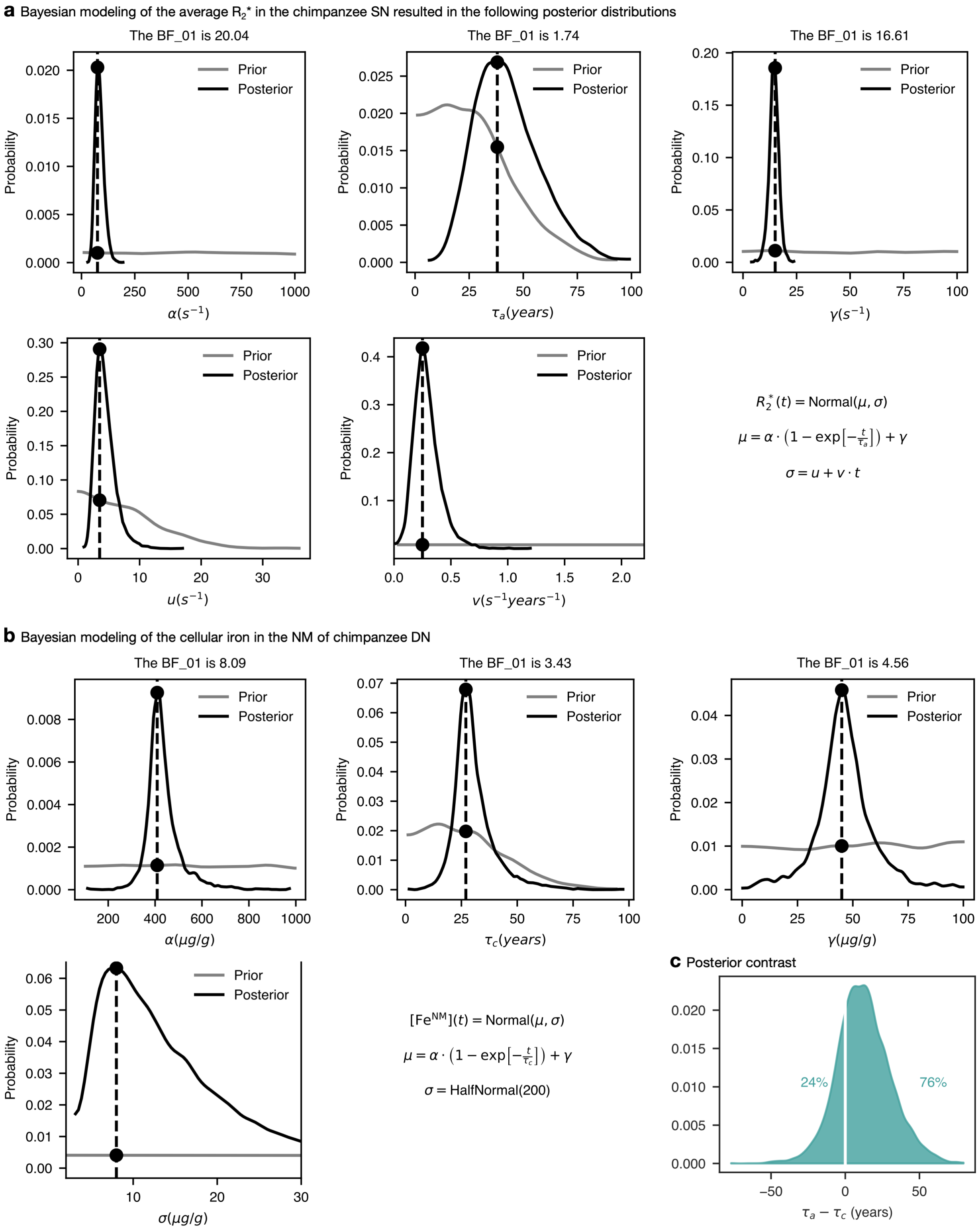


**Figure 12:** Prior and posterior distributions for the model parameters with indicated Bayes factors (BF) for **a** the average $R_2^*$ in the SN and **b** the cellular iron in the NM of chimpanzee DN. **c** To compare the lifespan dynamics of the iron accumulation in the DN and the average SN tissue, we calculated the difference in the posterior distributions of the time constants of iron accumulation in the DN, $\tau_c$, and $R_2^*$ in the SN, $\tau_a$. We conclude that the time constant for the iron accumulation in the DN is smaller than that for the iron accumulation in the surrounding tissue, as the posterior contrast $\tau_a - \tau_c$ is 76% positive, see supplement. On average, the time constant of iron accumulation in the SN measured by MRI is 13 years larger than for the cellular iron concentrations in DN (mean of the posterior contrast).

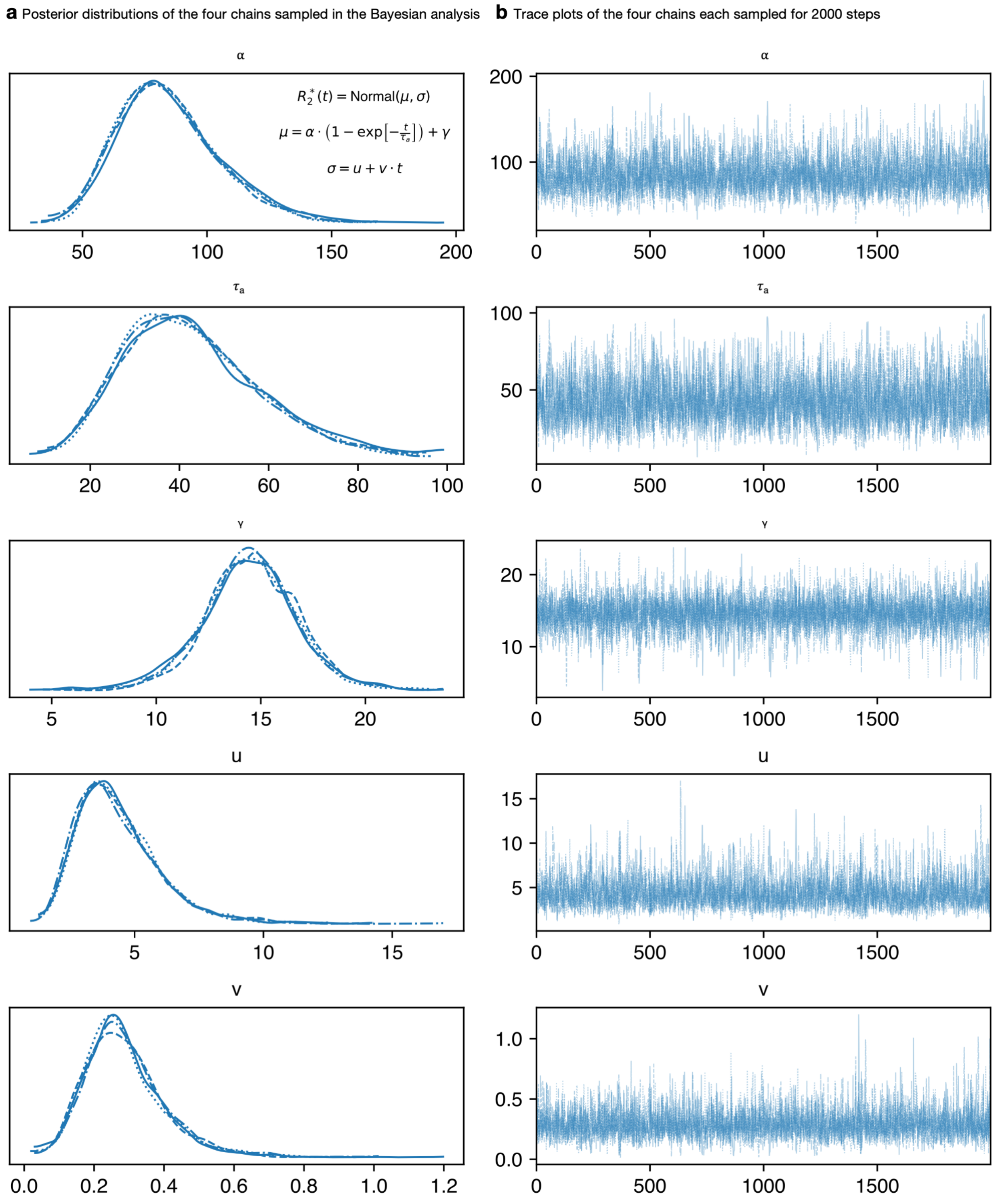


**Figure 13:** **a** Bayesian modeling of the average $R_2^*$ in the chimpanzee SN resulted in the posterior distributions for the five model parameters sampled by Markov Chain Monte Carlo. **b** The sampling processes of the four chains were monitored for quality assurance.

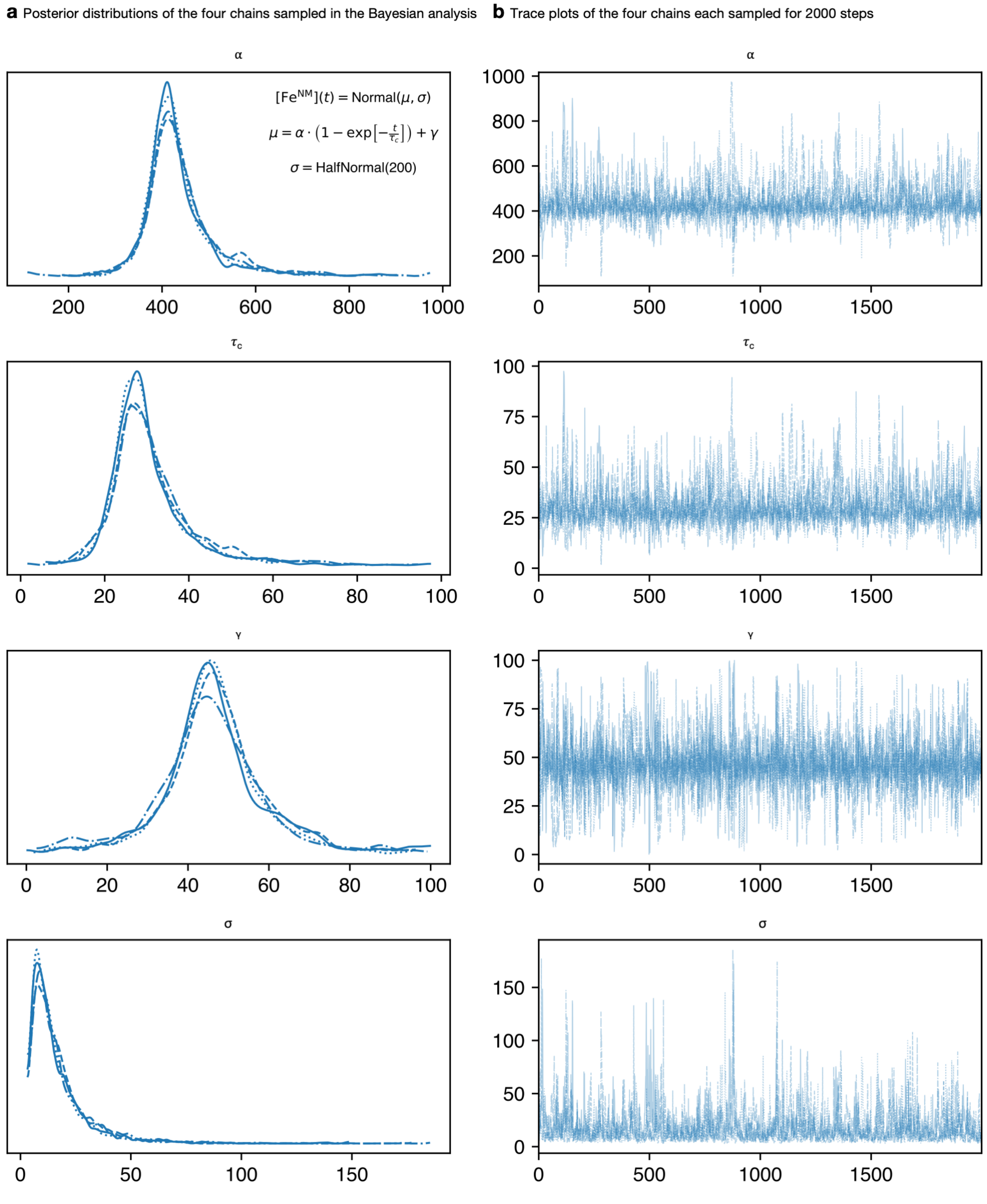


**Figure 14: a** Bayesian modeling of the cellular iron concentration in NM of chimpanzee DN resulted in the posterior distributions for the five model parameters sampled by Markov Chain Monte Carlo. **b** The sampling processes of the four chains were monitored for quality assurance.

#### 9.7 Iron binding dynamics

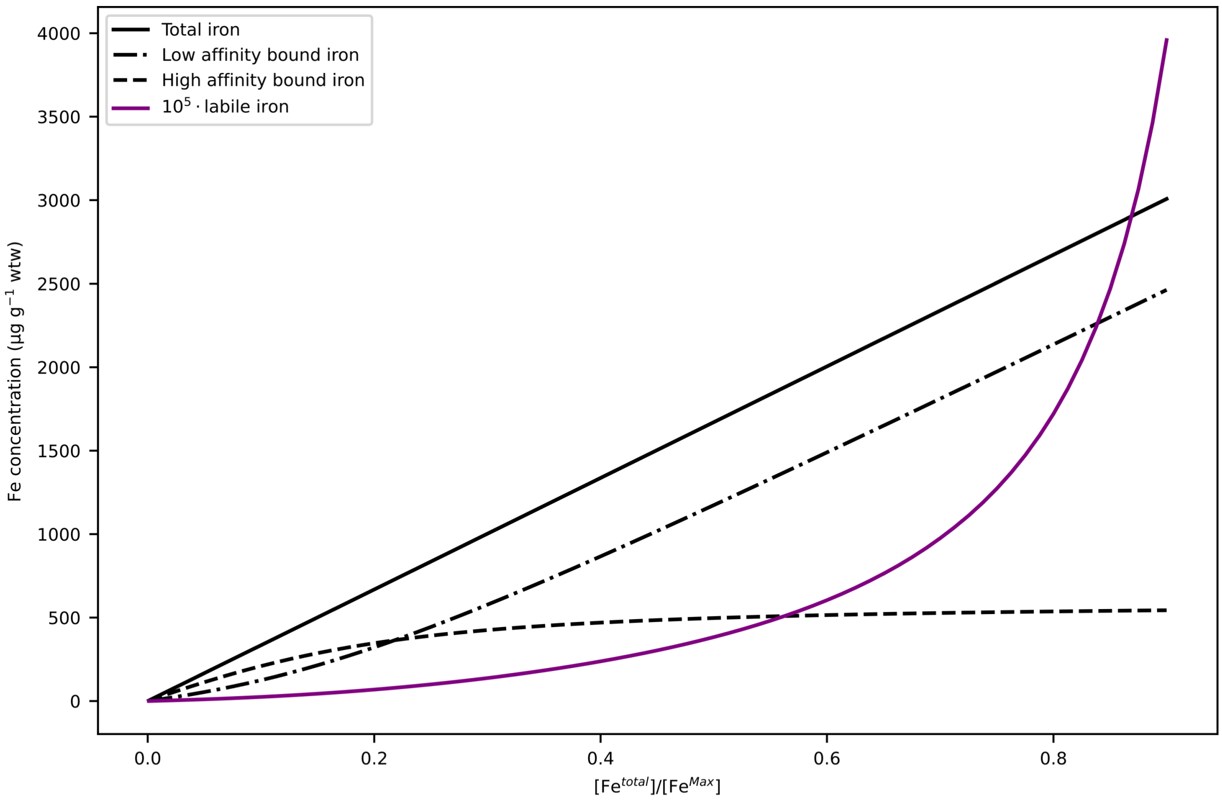


**Figure 15:** Dependence of the concentrations of labile iron (violet line, scaled by factor $10^5$ for visibility) and iron stored in the low- and high-affinity binding sites of NM (dashed and dotted lines) as a function of the parameter $\theta = \frac{[\text{Fe}^{\text{total}}]}{N \cdot H + N \cdot L}$. This shows that the labile iron concentration dramatically increases if both the NM-iron-binding-sites are saturated.
